# Time resolution in cryo-EM using a novel PDMS-based microfluidic chip assembly and its application to the study of HflX-mediated ribosome recycling

**DOI:** 10.1101/2023.01.25.525430

**Authors:** Sayan Bhattacharjee, Xiangsong Feng, Suvrajit Maji, Prikshat Dadhwal, Zhening Zhang, Zuben P. Brown, Joachim Frank

**Author notes:** These authors contributed equally (Sayan Bhattacharjee, Xiangsong Feng).

## Abstract

The rapid kinetics of biological processes and associated short-lived conformational changes pose a significant challenge in attempts to structurally visualize biomolecules during a reaction in real time. Conventionally, on-pathway intermediates have been trapped using chemical modifications or reduced temperature, giving limited insights. Here we introduce a novel time-resolved cryo-EM method using a reusable PDMS-based microfluidic chip assembly with high reactant mixing efficiency. Coating of PDMS walls with SiO2 virtually eliminates non-specific sample adsorption and ensures maintenance of the stoichiometry of the reaction, rendering it highly reproducible. In an operating range from 10 to 1000 ms, the device allows us to follow in vitro reactions of biological molecules at resolution levels in the range of 3 Å. By employing this method, we show for the first time the mechanism of progressive HlfX-mediated splitting of the 70S *E. coli* ribosome in the presence of the GTP, via capture of three high-resolution reaction intermediates within 140 ms.

## Introduction

To comprehend the fundamentals of any biological process, one requires insights into the underlying molecular mechanisms. It is often possible to study a reaction in vitro, outside the context of the cell. However, interactions among reactants and concurrent conformational changes of the molecules are too fast to be captured structurally using standard methods of biophysical imaging. In single-particle cryo-EM, the conventional pipetting-blotting method of sample deposition on the grid requires several seconds at least -- too long to capture reaction intermediates of molecular machines, which are in the range of tens or hundreds of milliseconds. In the past, some of such intermediates have been trapped by the use of non-hydrolysable analogs (such as GMP-PNP^1,2^, antibiotics^3^, or cross-linking^4^), but the insights gained in this way are limited and often burdened with presumptions.

Time-resolved cryo-EM (TRCEM) opens a way for obtaining structural and kinetic information on reaction systems that are not in equilibrium but change over time until equilibrium is reached^5^. In TRCEM studies, the biological reaction is started by mixing reactants, then stopped at a selected time point by fast-freezing, and the so trapped reaction intermediates are visualized by single-particle cryo-EM. By means of multiple experiments with different time points, TRCEM is able to capture the time course of a reaction, leading to a ‘movie’ of intermediates on the path to the state in equilibrium^6^.

Over the years, a variety of TRCEM methods have been developed^7-19^ showing the potential for obtaining key insights into the mechanism of action of molecules and molecular machines on the time scale of tens to hundreds of milliseconds. These methods can be divided into two categories: spraying/mixing^15^ and mixing/spraying^10^. In the first category, a specialized sprayer^15^ or dispenser system^7^ is employed to deposit one reactant onto a grid already covered with another reactant. Both mixing and reacting occur on the grid immediately prior to vitrification. Questions have been raised over the uniformity of a diffusion-dependent reaction on the grid^6^. In addition, electron tomography has shown that molecules are frequently observed to congregate at the top and bottom of the ice layer^20^, raising concerns about artifacts stemming from their extended exposure to the air-water interface in spraying/mixing or dispensing/mixing methods. In the second category, a microfluidic chip is used for mixing and reacting two reactants and for subsequent rapid spraying to deposit the reaction product onto the grid, followed immediately by plunging of the grid into the cryogen^8-10,12-14,16^. In that case, mixing and reacting can be efficiently controlled, and the time during which the reaction product is exposed to the air-water interface is kept minimal.

There are still some key issues among the existing methods which need to be resolved. One problem is sample adsorption on the walls of the chip. Polymers such as PDMS, IP-S, IP-Q are now commonly used as a cost-effective material to fabricate microfluidic chips^12,13,18,19^, but as a rule, the surfaces of these materials are intrinsically hydrophobic and adsorb a substantial amount of protein. In this case, the contact between the sample and the polymer microchannel can degrade the quality of the reaction, casting doubt over the accuracy of the kinetic information. This problem is avoided with silicon-based microfluidic chips^16,17^ which are by nature hydrophilic, but they are quite impractical as it takes several weeks and on average hundreds of dollars to manufacture a chip of a specific design (i.e., in multiple copies; with production steps including the etching and dicing of a silicon wafer and bonding it with glass). Another problem is insufficient initiation of a reaction, which may result from ineffective mixing of fluids due to limited micromixer performance^12,14^ in the laminar flow regime.

In the following we describe our TRCEM setup utilizing a PDMS-based microfluidics chip assembly of new design that overcomes these problems, as it efficiently mixes the reactants for uniform initiation of a reaction, conducts a controlled reaction virtually unimpeded by protein adsorption, and sprays the reaction product in a uniform three-dimensional cone onto the EM grid. We have used this novel device successfully in several studies on translation. Here we present the study of HflX-mediated recycling in detail to demonstrate the efficacy of the novel TR device, and its capability to yield biologically significant information.

**H**igh **f**requency of **l**ysogenization **X** (HflX) is a universally conserved protein for prokaryotes, a GTPase which acts as a ribosome-splitting factor in response to heat shock or antibiotics^2,21-23^. *E. coli* HflX consists of four domains: N-terminal domain (NTD), GTP binding domain (GBD), C-terminal domain (CTD), and helical linker domain (HLD)^2,24^. The cryo-EM structure of the HflX-50S complex stalled in the presence of GMP-PNP, a non-hydrolysable GTP analog, revealed that the HLD and NTD of HflX bind to the peptidyl-transferase center, presumably causing rupture of the intersubunit bridge B2a (h44:H69), thereby promoting the dissociation of the 70S ribosome^2^. However, the molecular mechanism of these events and particularly the interaction between HflX and 70S have remained elusive. Using TRCEM, we were able to capture three short-lived intermediate states, at resolutions in the 3-Å range, by starting the reaction between HflX and the *E. coli* 70S ribosome in the presence of GTP and stopping it at 10, 25 and 140 ms. Atomic models of these states allowed us to elucidate the mechanism of this process in great detail.

## Results

### A novel microfluidic chip assembly for TRCEM

The complete setup for TRCEM grid preparation, originally based on an apparatus built by Howard White^25^, is depicted in Figure S2. The heart of the TR apparatus is the microfluidic chip assembly mounted next to the plunger (Figure 1A), which are both accommodated in an environmental chamber that maintains the temperature and humidity at controllable levels. The plunger, which is pneumatically operated, holds the tweezers on which the EM grid is mounted for fast plunging into liquid ethane after passing the spray cone. In addition, the apparatus contains the pumping system for introducing the solutions into the micro-mixer and the nitrogen gas into the gas inlets of the micro-sprayer. Finally it also houses the computer for controlling both the pumping system and the plunger.

**Figure 1.**
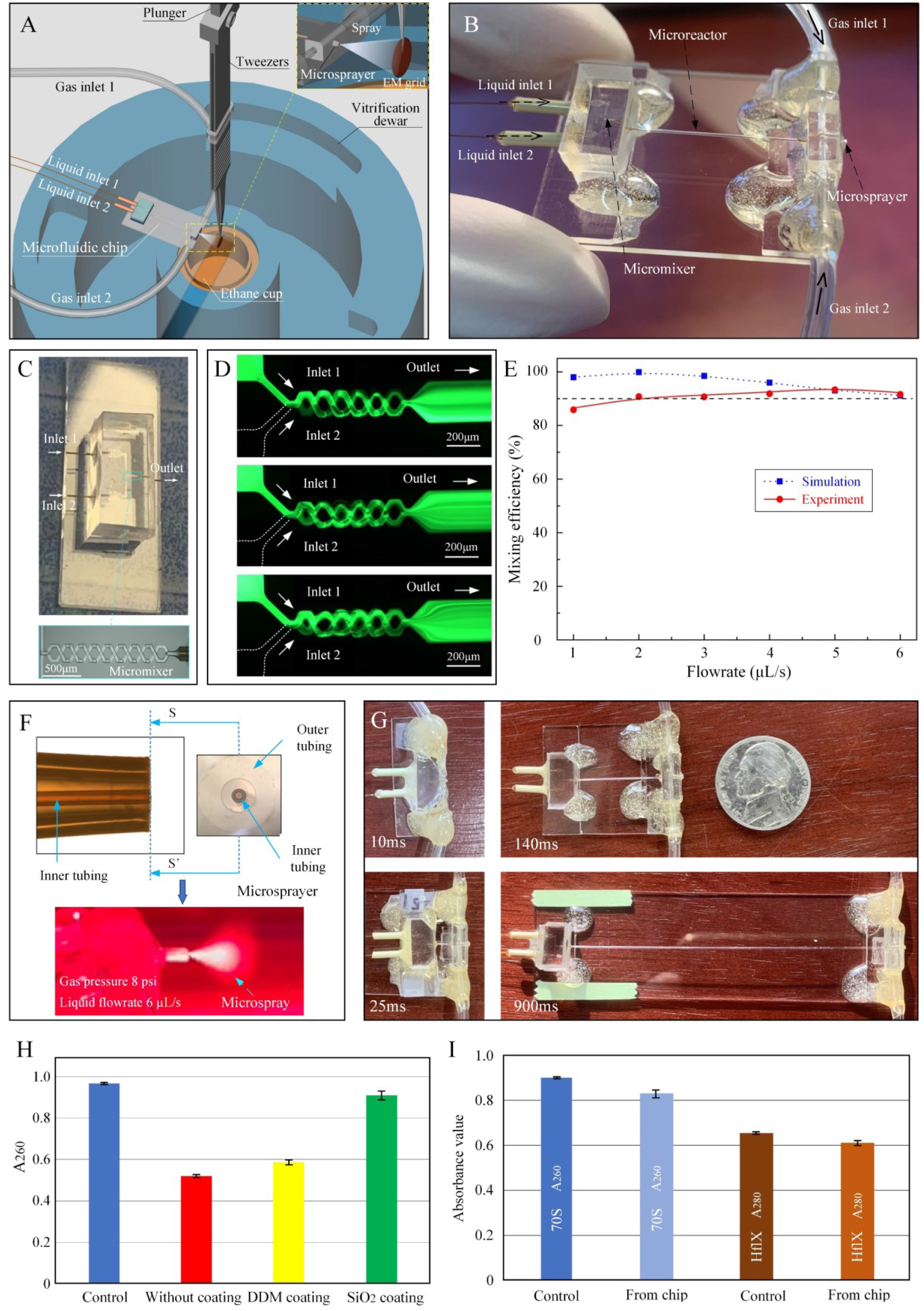
The modular PDMS-based microfluidic chip assembly for TRCEM sample preparation. (A) Schematic showing the setup for TRCEM using the mixing-spraying-plunging method. (B) The microfluidic chip assembly comprises three parts: micro-mixer, micro-reactor and micro-sprayer. (C) The splitting-and-recombination (SAR) based micro-mixer is fabricated by soft-lithography. (D) Fluorescence distribution along the micromixer with five mixing units under different inlet flow rate conditions. The mixing efficiency is characterized by the evenness in the distribution of fluorescent fluid. (E) Mixing efficiency for the micro-mixer at the exit under different flow rate conditions. The high mixing performance of this micro-mixer was validated both numerically and experimentally. (F) Micro-sprayer, with inner and outer tubing aligned and centered, used for depositing the reaction product onto the grid. The micro-spray (illuminated by red laser) is generated under the conditions of liquid flow rate 6 µL/s and gas pressure 8 psi, (G) A set of microfluidic chips was employed to achieve required reaction time points of 10, 25, 140, and 900 ms for the HflX study. (H) Compared with PDMS surface without any coating layer, and with DDM coating, the SiO_2_ coating shows effective mitigation of protein adsorption (*E. coli* 70S is used as sample). (I) The SiO_2_ coating functions well even after one month (here both *E. coli* 70S and HflX are used as samples).

We designed and successfully tested the modular TR chip assembly shown in Figure 1B, which is composed of three elements/modules: 1) a SiO2-coated, PDMS-based splitting-and-recombination (SAR) micro-mixer with 3D self-crossing channels (Figure 1C), which is able to mix the solutions with the effectiveness of > 90% at working flowrate of 6 μL/s (Figures 1E and S3, and Methods section in Supplemental Information(SI)); 2) a micro-capillary glass tubing serving as the micro-reactor for stable (i.e., unchanged under conditions of high pressure drop) reaction time control (Figure S4, and Methods section in SI); 3) a PDMS-based micro-sprayer with inner capillary tubing for spraying out the reaction product under the action of pressurized nitrogen gas (Figure 1F, Methods section in SI, and supplemental video 1).

**Figure 2.**
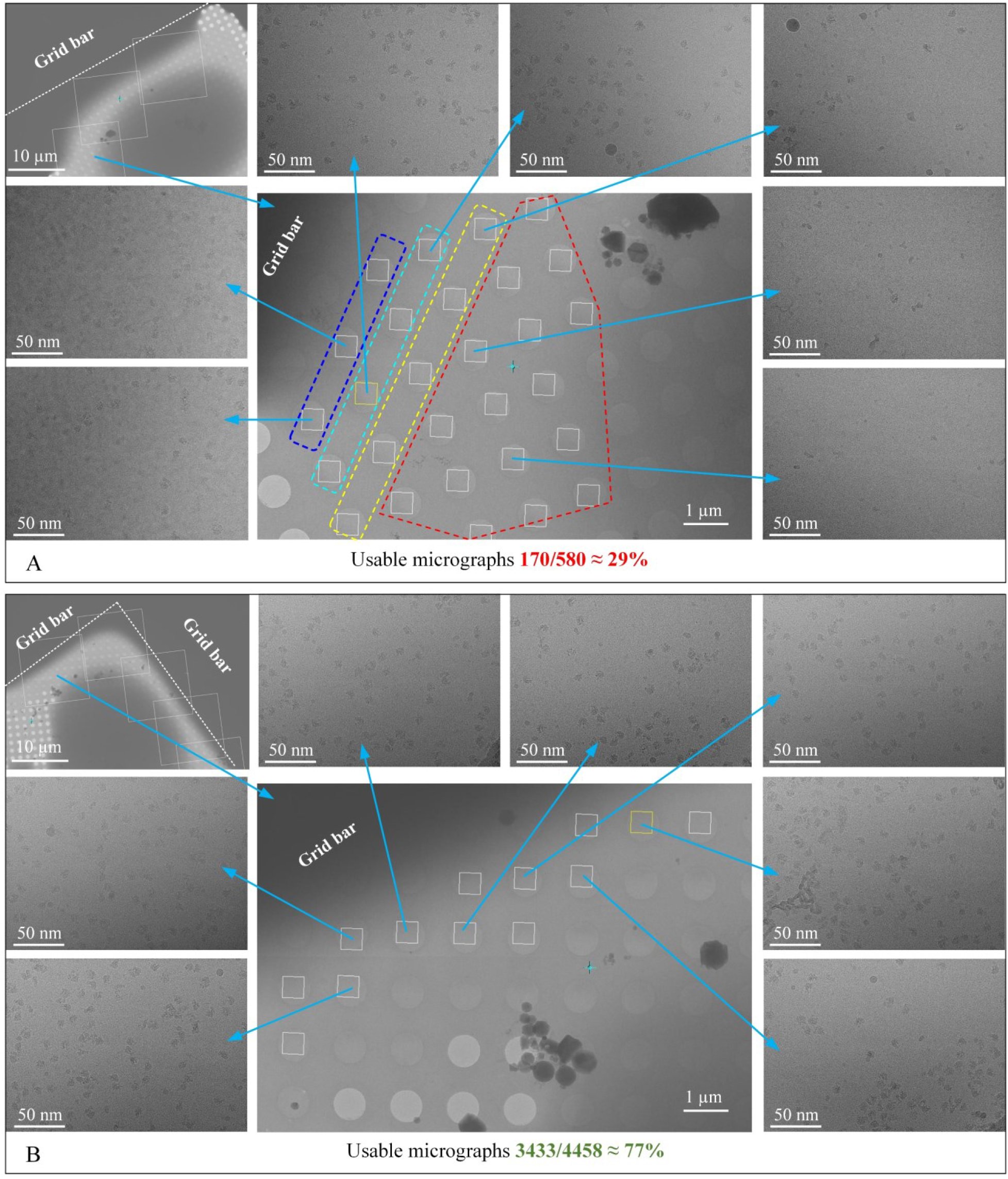
Data collection strategies on droplet-based cryo-grid prepared by mixing-spraying TRCEM method. (A) Collect as many micrographs as possible on each droplet. The number of particles gradually decreases when the target moves away from the grid bar, in the direction of areas marked in blue, cyan, yellow and red. (B) Collect only along two or three lines of holes which are near and parallel to the grid bar.

**Figure 3.**
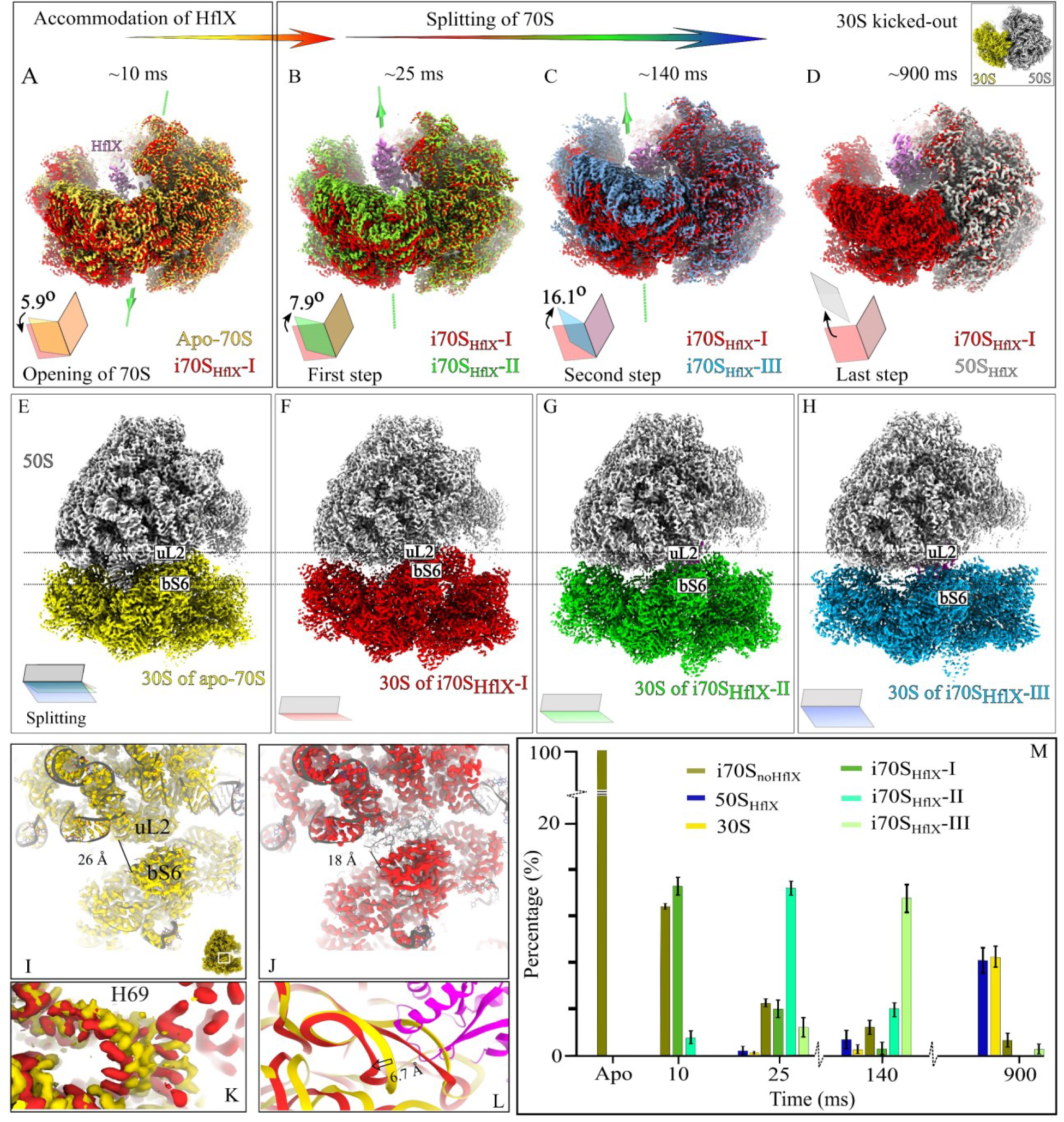
Molecular details of subunit interface during the progressive opening of the 70S. **(A)** Superimposition of reconstructions to show the opening of the 30S subunit from apo-70S (yellow) to i70S_HflX_-I (red) to accommodate HflX. The green line represents the initial axis of 30S rotation, Axis I. (**B**) and (**C**), reconstructions of second and third intermediates overlapped with first intermediate, showing the stepwise splitting of the 70S by HflX by rotation of the 30S subunit around Axis II (green line). The corresponding rotation angles and direction of 30S rotation are shown in cartoon book representations. (**D**), reconstruction of the 50S-HflX complex after the departure of the 30S subunit, overlapped with the first 70S intermediate. In (**A**) through (**D**), all reconstructions are aligned on the 50S subunit. (**E**) to (**H**) are the high-resolution densities of control 70S at 900 ms and three intermediates obtained within 140 ms, and showing the stepwise rotation of 30S during recycling with respect to 50S (gray). (**I**) and (**J**) are the zoomed views of Coulomb densities in yellow for apo-70S and red for i70S_HflX_-I, respectively, and corresponding atomic models (gray) showing the rearrangement of the protein uL2 of the 50S and bS6 of 30S to accommodate HflX. (**K**) and (**L**), Coulomb densities, and corresponding ribbon models of H69 from apo-70S (yellow), and i70S_HflX_-I (red), respectively, showing the very first movement of helix H69. HflX is shown in magenta. (**M**) Kinetics of the splitting reaction in terms of the number of particles per class as a function of time, obtained by 3D classification.

To prevent adsorption of molecules, plasma-enhanced chemical vapor deposition (PECVD) is used to coat the inside walls of the PDMS micro-mixer channels with a thin SiO2 layer. We tested the sample adsorption with the *E. coli* 70S ribosome to compare the chips without coating to those with DDM or SiO2 coating. The sample concentration in buffer is measured before and after passing the devices with different types of coatings by our Nanodrop UV-Vis Spectrophotometer (see Methods section in SI). In this experiment, 94% of the initial concentration was retained using the SiO2-coated chip (Figure 1H and Table S2), while only 54% and 60% were retained after the sample was passed through the chip assembly without coating or with DDM coating, respectively. These findings demonstrate that the SiO2 coating can effectively mitigate the sample adsorption for ribosomes. After one month, the SiO2-coated chip assembly was tested again with 70S ribosome and HflX protein, and 92% of the 70S and 93% of HflX were shown to be still retained, respectively (Figure 1I). These results demonstrate that the hydrophilicity of the internal surface is virtually undiminished after a substantial period of time.

Thus it is apparent that the solutions introduced from the glass-capillary liquid inlets will pass through the entire microfluidic device (SiO2-coated micromixer, glass-capillary micro-reactor and glass-capillary inner tubing of the micro-sprayer) without contact with any hydrophobic surface, a fact of high importance for preserving the stoichiometry of a reaction and guaranteeing its reproducibility.

Based on these initial test results, we fabricated a set of microfluidic chips (Figure 1G) to conduct a set of biological experiments. The relevant materials and parameters for the fabrication of the chip assemblies are listed in Table S1. An estimation of reaction times achieved using this TRCEM method in the application to HflX-mediated ribosome recycling is given in the Methods section in SI. Four microfluidic chip assemblies were used, with reaction times estimated to be 10, 25, 140, and 900 ms, respectively.

### Exposure-targeting strategy in data collection for droplets-based cryo-grids

For grids obtained by the conventional blotting method, data collection is usually done automatically after the template is set up for targeting both the holes and exposures. But for grids prepared by the mixing-spraying method, it is not easy to use automation, since every collectible square possesses droplets of different sizes and thicknesses^25^. Hence time-consuming manual exposure targeting is required.

Some observations on typical particle distributions in the HflX experiment, to be detailed below, and other experiments led us to develop an effective strategy for exposure targeting in data collection on droplet-based grids.

There are two types of situations: one is where the droplet has no contact with the grid bar (marked red in Figure S8); and the other where it does have contact with the grid bar (marked green in Figure S8). In the former, the ice is observed to be thick and unsuitable for data collection, while in the latter, there is always some part of the region near the grid bar with an ice thickness suitable for data collection. All our exposure targets are therefore focused on the second type of droplets, as detailed below.

In the beginning, as shown in Figure 2A, we tried to collect data on as many holes as possible for droplets of the second category, i.e., touching or partially covering the grid bar, and found that typically there are four regions with different behaviors, following a trend: (i) very close to the grid bar, as in the area marked blue, the ice is not vitrified very well but particles are still visible; (ii) in the area marked cyan, particles are clearly visible; (iii) further on, in the area marked yellow, the number of the particles has significantly decreased; until, (iv) in the area marked red, there are almost no particles left (Figure 2A). In the present instance of data collection, only 170 out of 580 micrographs, or 29%, were left for data processing.

In line with these observations, we developed the exposure-targeting strategy shown in Figure 2B: we collect only along two or three lines of holes which are near and parallel to the grid bar, in the areas of type (i), (ii) and (iii). As a result, in our example, we obtained 3433 good micrographs out of a 4458 total, which means about 77% could be used in this case for data processing. We therefore adopted this strategy for all our data collection on grids prepared by TR cryo-EM.

### Time-resolved experiments on HflX-mediated ribosome recycling from 10 to 900 ms

HflX acts on the 70S ribosome in a nucleotide-dependent way, and light scattering analysis revealed that the rate of ribosome splitting by HflX-GTP (0.002 s^-1^) is very similar to the rate of ribosome dissociation by the combined action of RRF and EF-G-GTP (0.005 s^-1^)^22,26^. The fraction of ribosomes split into subunits at room temperature within a reaction time of 140 ms is close to 50%, according to our earlier TR cryo-EM experiment on *E. coli* ribosome recycling in the presence of RRF, EF-G, and GTP^9^. In view of these findings, we anchored our TR cryo-EM study at a 140 ms reaction time point and added two shorter time points (10 ms and 25 ms) and one longer one (900 ms) toward the reaction’s completion. We mixed 70S ribosomes with the HflX-GTP complex in our mixing-spraying TRCEM apparatus (Figure S2) using different microfluidic chips of the PDMS-based design (Figure 1G and Methods section in SI). As in our previous TRCEM studies^9,10^, 3D classification was performed on the entire, pooled dataset across all four time points. The 3D classification produced seven distinct classes, which we characterized by examining the corresponding reconstructed density maps (note: “rotated” and “nonrotated” refers in the following to the presence or absence of intersubunit rotation^27^): (1) rotated 70S without HflX (r70SnoHflX); (2) non-rotated 70S without HflX (nr70SnoHflX); (3) 70S-like intermediate-I with HflX (i70SHflX-I); (4) 70S like-intermediate-II with HflX (i70SHflX-II); (5) 70S-like intermediate-III with HflX (i70SHflX-III); (6) 50S with HflX (50SHflX); and (7) 30S (Methods section in SI and Figures S10, and S11).

The splitting reaction kinetics of the 70S ribosome, as evaluated by counting the numbers of particles obtained upon 3D classifications from 10 ms to 900 ms, is found to follow a similar, roughly exponential behavior as reported from dissociation kinetics measured by light scattering^2^ (Figure 3M). Furthermore, we noticed a rapid increase in the number of free 30S subunit particles from 140 ms to 900 ms, which leads us to conclude that the final separation of the subunits commences not earlier than with state i70SHflX-III (Figure 3M).

### Intermediate states of HflX-mediated recycling and their interpretation

The three classes of HflX-containing intermediates and class 50SHflX -- four of the seven 3D classes we found -- were selected for additional structural analysis (Methods section in SI and Figures S10, and S11A-E). Furthermore, focused 3D classification and subsequent reconstruction of HflX-binding regions from each of the resulting class reconstructions yielded high-resolution density maps for four states of HflX: (1) HflX-I, (2) HflX-II, (3) HflX-III, and (4) HflX-IV (Figure S10). Refinement on the three i70SHflX class reconstructions yielded high-resolution on-pathway intermediates i70SHflX-I, i70SHflX-II, and i70SHflX-III (Figures 3A-H), and their resolutions are indicated in Figure S12 and Table S3. The kinetics of the reaction can be followed from the histogram of particle counts in the respective classes (Figure 3M). The intermediates i70SHflX-I, i70SHflX-II, and i70SHflX-III are each dominant in the 10 ms, 25 ms, and 140 ms time points, respectively, but are always intermixed with the other intermediates, as well as with apo-70S, and the 50S-HflX end product.

In all three intermediates, the CTD of HflX is found anchored to uL11 of the 50S subunit at the bL12 stalk base. Overall the comparison of the intermediates shows a gradual clamshell-like opening of the 70S ribosome (see supplemental video 2).

Structurally, the intermediates are distinguished by (i) the degree of the clamshell-like opening, (ii) the position of helix H69 (in two steps, from i70SHflX-I to -III), (iii) the position of helix H71 (from i70SHflX-I to -II), and (iv) the position of HflX with respect to the 50S subunit (from i70SHflX-I to -II, and reversed from -II to -III). In the final state observed, after the departure of the 30S subunit, HflX remains bound to the 50S subunit.

Comparison of the atomic models obtained for these intermediates with one another and with the apo-70S revealed that the opening and splitting of the 70S ribosome occurs in the following steps:

First, the ribosome opens slightly to accommodate the initial binding of HflX in i70SHflX-I (Figures 3A, 3E-F). Using the tool previously developed^28^ we find that in this first intermediate, the 30S subunit has rotated by 5.9° around an axis (Axis I) that passes through the intersubunit bridges B1b, B2a, B3, and B4 (Figures 6A, D, G, and 6J-K), and this rotation has moved protein bS6 of 30S into close vicinity to protein uL2 of the 50S subunit (Figures 3I-J). Apparently, the insertion of HflX along with the prying apart of the 70S ribosome and the rotation of the 30S subunit is driven by the increase in backbone entropy of uL2 in i70SHflX-I compared to apo-70S since we find indications of disorder: the density of uL2 is not resolved well in i70SHflX-I (Figure 3J) compared to all its other manifestations in apo-70S, i70SHflX-II and i70SHflX-III (Figures 3I-J, and S13). Comparison of the 50S subunit in i70SHflX-I and apo-70S shows that H69 has moved by 6.7 Å, apparently through a push by HflX since fitting the model of HflX to apo-70S reveals a steric clash with H69 (Figure 3K-L). In i70SHflX-I HflX is blurred, indicating motion-induced heterogeneity of the population in that class (Figure S10).

Going from this first intermediate to i70SHflX-II and i70SHflX-III we observe stepwise rotations, by 7.9° and 8.2°, respectively, of the 30S subunit around a new axis (Axis II) passing through intersubunit bridges B3 and B7a, which are both located along helix h44 (Figures 3B-C, 3F-H, and 6B-C, 6E-F, 6H-I, and 6J-K). In the first step of rotation around this new axis, protein bS6 moves away from protein uL2 (Figures 3B, 3F-G, and 6B, 6E, 6H, and 6J-K). This movement is made possible by a 6.5-Å pull of C1965 of H71 as a result of HflX moving from its previous position on i70SHflX-I to a new position on i70SHflX-II and a subsequent shift of the loop-helix motif (G74-V100) associated with the NTD of HflX (Figures 4D-G). As a consequence, bridge B3 (h44:H71), as well as bridges B7b and B7bR, have become destabilized. While the conformation of the 30S subunit remains virtually the same from apo-70S to the first intermediate, the change from the first to second intermediate is accompanied by a rotation of the 30S subunit head by 2.1° around another axis, Axis III (Figure S14).

**Figure 4.**
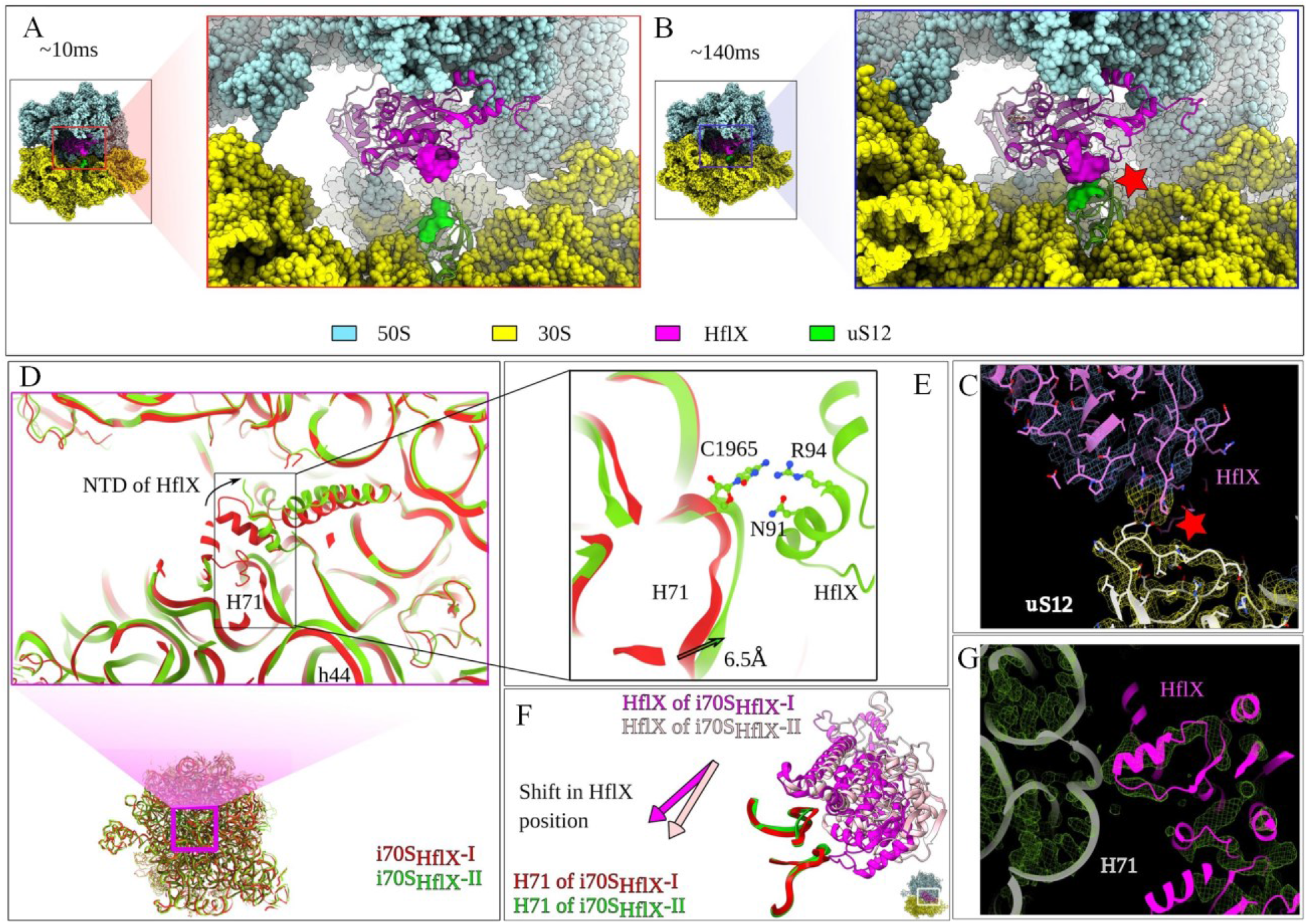
The shift of S12 towards HflX and Involvement of HflX in 70S splitting. The zoomed view of HflX and S12 of 30S interactions from i70SHflX-I and i70SHflX-III states are shown in (**A**) and (**B**), respectively. Due to the stepwise separation of the 30S during splitting, there occurs a steric clash (red star) at 140 ms due to the subsequent shift of the whole S12, which is not present at earlier states like10 ms, and this clash is the cause of the final separation of 30S from 50S by a power-stroke from HflX upon GTP hydrolysis. (**C**) The same clash at 140 ms from i70SHflX-III is shown in Coulomb density with a fitted model. (**D**) Pulling of H71 by the NTD of HflX causes the disruption of intersubunit bridge B3 between H71 and h44, and the zoomed view in (**E**) shows the interacting residues of both H71 and HflX. During pulling of H71 HflX has to shift its position and is shown in (**F**). (**G**) The same interaction is shown in Coulomb density with a fitted model.

In the second step of the 30S subunit rotation around Axis II, from i70SHflX-II to i70SHflX-III, protein bS6 has continued to move away from protein uL2 (Figure 3C, 3F-H, and 6C, 6F, 6I, and 6J-K), and HflX has shifted back to its original position on the 70S ribosome in i70SHflX-I. Bridges B7b and B7bR are now entirely disrupted, allowing the flexible loop (E323-G349) of HflX’s GTD to readily access the 30S subunit protein uS12, thus positioned to jettison the 30S subunit from the 70S ribosome (Figure 4A-C).

Finally, the reconstruction of the stable 50S-HflX complex, at 3.6-Å resolution (Figures 3D, and S15A), no longer shows any trace of density from the 30S subunit (Figures S15A, and D). This class mainly contains particles from 900 ms (Figure 3M). The map agrees quite well with the map of HflX-50S-GNP-PNP^2^ (Figures S15A-C).

A comparison of the atomic models built for the three intermediate states reveals that HflX changes its position on the ribosome and undergoes major conformational changes, specifically in its CTD, HLD, and NTD. The domain movements of CTD and HLD match quite well with the dynamics of apo-HflX predicted from 1000 ns of molecular dynamics simulations (Methods section in SI and Figure S16). Interestingly, the loop-helix motif (G74-V100) of NTD makes stable contact with H71 of the 50S subunit in i70SHflX-II (Figures 4D-F).

### Dynamics of HflX and possible GTP-bound state

In trying to understand the actions of HflX, we examined the hydrolyzation state of GTP in the different states of HflX. At 25 ms, with the exception of their NTDs, the densities of HflX and associated nucleotides in states HflX-I and HflX-II are not resolved as well as they are for the other two states, indicating mobility and preventing determination of the hydrolyzation state (Figure S10). In an attempt to fit the atomic model of GTP to the corresponding densities in HflX-III and HflX-IV, we observed that the density in the region of the nucleotide site on HflX-III is a better fit for GDP·Pi than for either GTP or GDP (Figure 5A). A similar matching effort resulted in a decent match of GTP to the density of HflX bound to the 50S subunit, even though the GDP state is expected to be found at this stage (Figure 5B).

**Figure 5.**
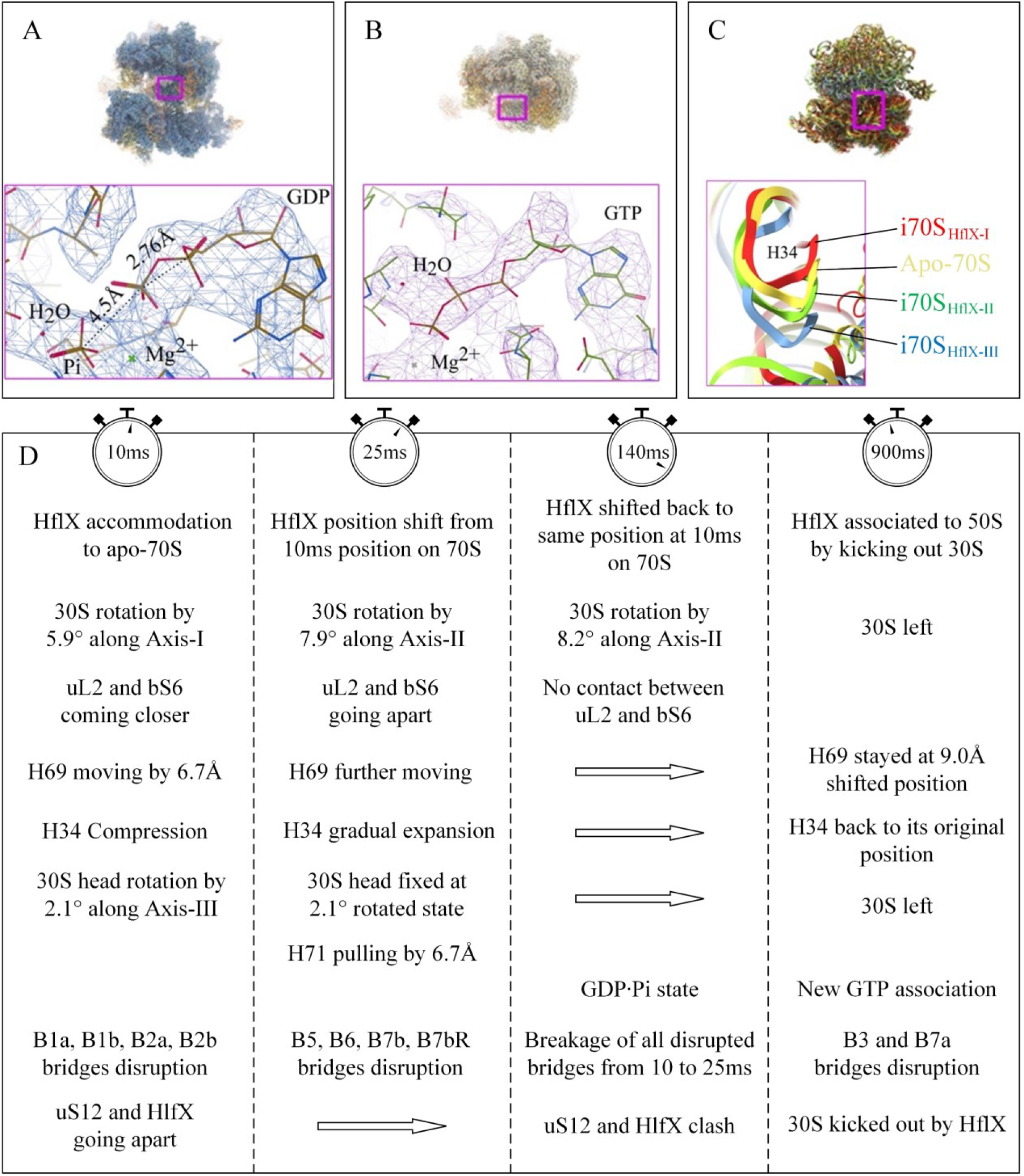
Analysis of nucleotide states, and spring-like nature of H34 and molecular events. **(A)** Refined density map of i70S_HflX_-III (in mesh) with an atomic model fitted in with the zoomed view of i70S_HflX_-III with the fitted model of GTP**·**Pi. The distance of 4.5 Å between GDP and Pi is compared with the distance of 2.8 Å between two P atoms of GDP. This distance between GDP and Pi is too large to form a P-O bond as in GTP. (**B**) Refined density map of 50S_HflX_ (in mesh) with atomic model fitted in along with the zoomed view of 50S_HflX_ with the fitted model of GTP. (**C**) The superimposition of atomic models of apo-70S and three intermediates with the zoomed view of H34 showing its spring-like behavior. Corresponding colors are indicated. (**D**) The molecular events involved during the 70S splitting by HflX are tabulated.

**Figure 6.**
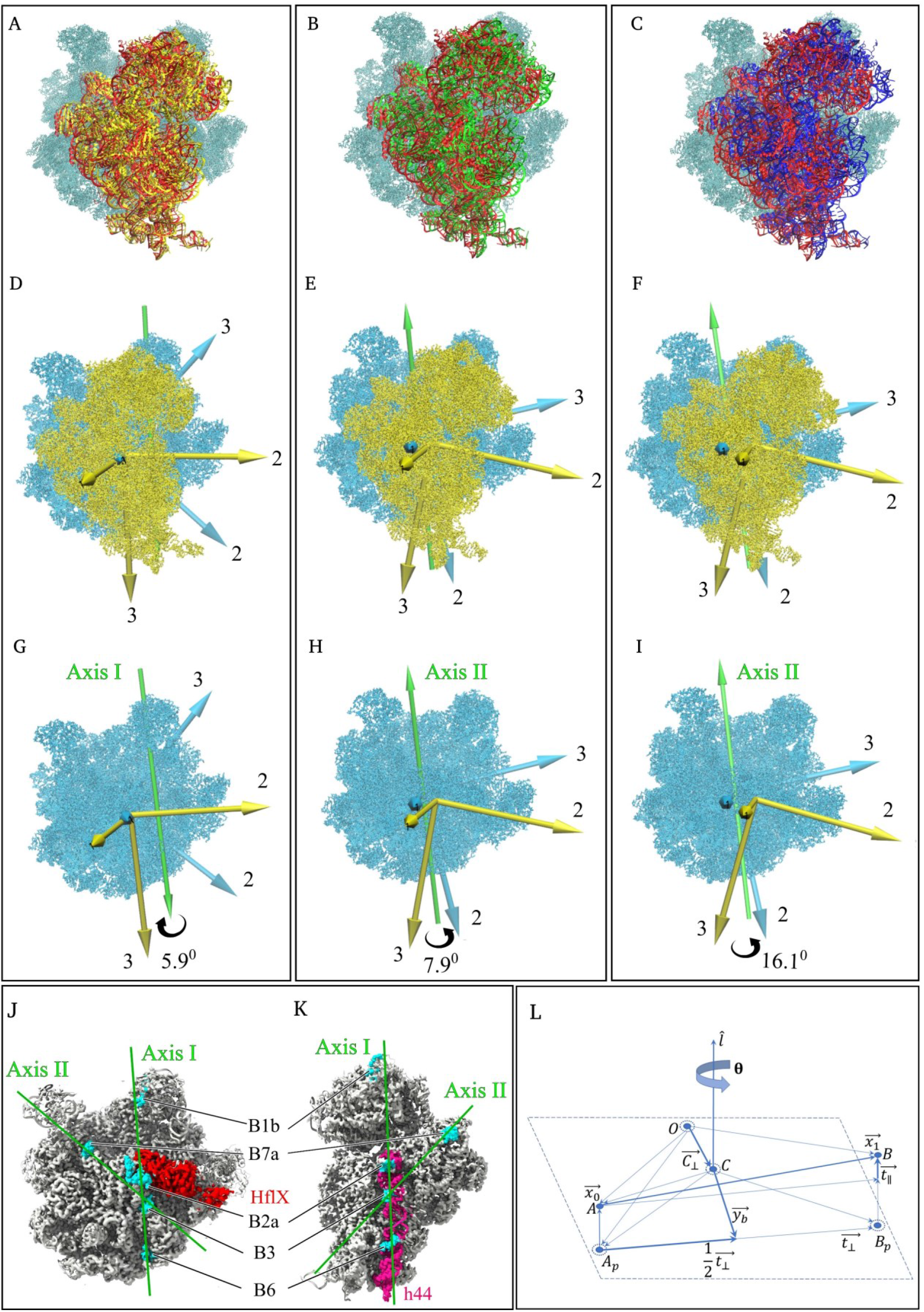
Axes of rotation of 30S subunit during splitting/recycling of the 70S ribosome: (**A**), (**B**), and (**C**), rotation of 30S subunit from Apo-70S (yellow) to i70S_HflX_-I (red), i70S_HflX_-I (red) to i70S_HflX_-II (green), and i70S_HflX_-I (red) to i70S_HflX_-III (blue), respectively. **(D)**, **(E)**, and **(F)** characterization of the 30S subunit rotation while the 50S subunits (cyan) are fixed in space. The models show the rotated state of the 30S subunit in each case. For clarity, 30S subunits are represented with only their principal axes of inertia. Rotation axes (Axis I, Axis II) are shown as green arrows indicating the direction (right-hand thumb rule) of the rotation (black curled arrows). **(D)** Rotation by 5.9° of 30S subunit around Axis I from Apo-70S (yellow) to i70S_HflX_-I (red). (**E**) and (**F**), Rotations by 7.9° and 16.1°, respectively, of 30S subunit around Axis II. **(G)**, **(H)**, and **(I)**, same representation as **(D)**, **(E)**, and **(F)** omitting the 30S subunit to show the axes clearly. (**J**) and (**K**), Intersubunit bridges through which Axes I and II pass on the 50S and 30S subunit, respectively. **(L)** Formulation of rigid body motion along with location of the rotation axis. Initial and transformed positions are respectively denoted by A (position vector *x→*_0_) and B (position vector *x→*_1_). The shift from A to B is given by translation vector *t→*. Points with dotted circles are on the plane perpendicular to the rotation axis *l*^^^. Points *A_p_→* and *B_p_→* are projections of points A and B, respectively. *t→*_∥_ and *t→*_⊥_ are respectively the parallel and perpendicular components of the translation vector *t→*. Axis *l*^^^ and angle θ describes the rotation of the rigid body from point A to B. The rotation axis *l*^^^ passes through the point given by the position vector *C→*_⊥_. The solution for the position vector *C→*_⊥_ is obtained by using the remaining vectors, as indicated in the derivation in Methods Section in SI “Determining the position vector of the unique point through which the rotation axis passes”.

Since at the 140 ms time point the inorganic phosphate (Pi) is still associated with GDP and virtually none of the 30S subunits has been cleaved off, we conclude that the energy for the breaking of bridges B3 and B7a and the final dissociation of the 30S subunit is set free by the release of Pi. It is unclear without further investigation if GTP hydrolysis plays a role in the initial stages of splitting, from intermediate I to II, since it is known that HflX can perform the splitting in the absence of GTP, albeit at a slower rate^2^. The likely explanation for the observation of GTP on the 50S subunit-bound HflX molecule at 900 ms is that by that time both Pi and GDP have left and that a new GTP molecule has taken their place.

### Time-dependent rupturing of intersubunit bridges

As the direct observation of the state of bridges from the cryo-EM map is not conclusive due to resolution limitations, we proceeded with a geometric calculation to estimate the sequence in which intersubunit bridges are ruptured. With the axes and angles of the 30S subunit rotation known, as well as the locations of all bridges relative to the axes, we were able to determine the distances between constituent residues of all intersubunit bridges. From these distances and known ranges of chemical bond lengths we were able to estimate at which time points the intersubunit bridges are likely disrupted and broken (Figure 5D). According to these calculations, bridges B1a, B1b, B2a, and B2b are already disrupted at 10 ms. Bridges B5, B6, B7b, and B7bR are disrupted between 10 and 25 ms. All these bridges are found to be broken within 140 ms. Finally, the last two bridges, B3 and B7a, which form the hinges of Axis II, give way at some time point between 140 ms and 900 ms. Bridge B4 (H34:uS15) presents an interesting case as it behaves like a spring: its 50S subunit constituent H34 is initially compressed in the step from apo-70S to i70SHflX-I, as HflX is accommodated within the first 10 ms, but in the next two steps (10 ms to 25 ms to 140 ms) it is extended (Figure 5C). This bridge finally breaks along with B3 and B7a after 140 ms and, in 50SHflX, helix H34 has returned to its original position as in apo-70S.

## Discussion

Here, we present a method for preparing time-resolved cryo-EM grids to capture intermediates on the ∼10 to 1,000-ms timescale. Different in design from those by Lu et al.^16,17^, Mäeotset al.^13^, and Kontziampasis et al.^14^, the microfluidic chip assembly comprises three replaceable modules: (1) a PDMS-based internally SiO2-coated micro-mixer of the splitting-and-recombination (SAR) type, for efficiently mixing two samples without significant problems from protein adsorption; (2) a glass capillary micro-reactor for defining the reaction time, and (3) a PDMS-based micro-sprayer for depositing the reaction product onto the EM-grid. The sample is subsequently vitrified by fast plunging of the grid into the cryogen so that the TR cryo-grid can be well prepared.

In our application to the study of HflX-mediated ribosome recycling, a bacterial defense response to stress, we have demonstrated how our method is able to capture, on-the-fly, high-resolution structures of intermediates that represent snapshots of an unfolding complex molecular mechanism. Based on our cumulative observations from the examination of three short-lived reaction intermediates, we propose that the initial binding of HflX within 10 ms is followed by a stepwise clamshell-like opening of the ribosome around an axis that closely aligns with helix 44 of the 30S subunit, and that rupture of the last two remaining intersubunit bridges, B3 and B7a, occurs after 140 ms and most probably as a result of Pi release.

### Further applications of time-resolved cryo-EM in the study of functional dynamics

Our study will spur interest in the community for extending these studies to recycling in the 80S ribosome by protein factors such as ABCE1 to structurally reveal an evolutionarily conserved mechanism. We believe, moreover, that the quantitative description of our results on a dynamic process – employing tensor analysis to determine the tilt axis and rotation of subunits, and quantifying the timing of intersubunit breakage – as well as the application of this microfluidic device will open a new direction in the characterization of molecular events by cryo-EM and will stimulate interest and draw considerable attention among a wide audience of structural biologists, microbiologists, and pharmacologists. Ultimately, it may help in the design of a new class of broad-spectrum antibiotics that are able to overcome antibiotic resistance.

## Limitations of the Study

### Limitation in time range

Our method of TRCEM has been proven powerful for capturing structural intermediates for HflX-mediated ribosome recycling and other processes of translation, but there are many other interesting biochemical processes awaiting visualization by this method. Those involving motions of large domains of macromolecules are excellent candidates. Signaling, activation and gating mechanisms of receptors and transporters^29-31^ come to mind, as well, but in many cases the characteristic times of motions are much shorter than 10 ms, the minimum reached by our method of mixing-spraying-plunging. In this time domain, entirely different technologies have to be considered^15,19,32,33^

### Limitation in the accuracy of kinetic information

As we pointed out before^10^, TRCEM, in addition to furnishing the high-resolution structures of reaction intermediates, has at least in principle the capacity to give kinetic information, as well, since the numbers of particles in each structural class are known. However, this information is currently not very accurate since it depends on the vagaries of particle picking and classification strategies. Investment in a search for quantitative, reproducible strategies would therefore be of enormous value.

### Inefficiency of data collection

Data collection on droplet-covered TRCEM grids remains quite time-consuming even with the proposed strategy to target certain regions close to the grid bar, since it still relies on visual/manual selection. There is clearly a need for sophisticated automatic targeting tools on grids prepared by spraying with sample droplets.

### Large required sample quantity

Compared with conventional blotting method, the TRCEM method requires a substantially bigger volume of sample (∼10 μL versus 3 μL) for each grid, which is the minimum quantity of fluid required to reside in the whole microfluidic system for ensuring a stabilized spray. However, much of the spray is wasted in the present setup with a single EM grid as target, and an obvious next step is the development of a plunger with multiple pairs of tweezers or a specially designed tweezer-manifold to hold several grids at once.

### Unresolved questions regarding the mechanism of HflX action

Our TRCEM analysis about the interaction between HflX and 70S ribosome leaves open the question of how HflX recognizes the stalled state of the ribosome. Here our observation of 30S subunit head rotation from apo-70S to i70SHflX-I may offer a clue. Puromysine-treated polysomes, having deacylated tRNA in the P-site, display greatly enhanced HflX splitting activity, and this state was proposed as the natural substrate for HflX^2^. In this state, the ribosome is known to undergo spontaneous intersubunit rotation^34^, which goes hand in hand with 30S subunit head ‘swivel’ rotation^35^. This would suggest that HflX initially binds to the ribosome in its rotated conformation and forces it into the unrotated conformation observed in i70SHflX-I, with residual 30S subunit head rotation still retained. Another similarity holds if our hypothesis of HflX binding to the rotated ribosome is correct since the latter is the substrate of RRF/EF-G-GTP binding, as well^36,37^. But to answer these questions more extensive studies with similar tools are required.

Supplemental video 1. Spraying and plunging during the TR experiment.

Supplemental video 2. The clamshell-like splitting of the ribosome from Intermediate I over Intermediates II and III, ending with the 50S subunit (shown) and the 30S subunit (not shown), the end products of the recycling process.

## Acknowledgements

This work was supported by a grant from the National Institutes of Health R35GM139453 (to J.F.). All data was collected at the Columbia University Cryo-Electron Microscopy Center (CEC). We thank Robert A. Grassucci, and Yen-Hong Kao for their help with the cryo-EM data collection. The microfluidic chips with SiO2 coating were fabricated in Nanofabrication clean room facility in Columbia University.

## Author contributions

S.B., X.F., and J.F. conceived the research; S.B., X.F., and J.F. designed the experiments; S.B. prepared the biological samples; X.F. designed the TR chips; X.F., and P.D., developed the chip; X.F., S.B., performed the TR cryo-EM experiments; S.B., X.F., and Z.Z. collected the Cryo-EM data; S.B. processed the data; S.M. calculated the subunit and domain motions and the axes of rotation; Z.P.B. helped S.B. with atomic model building and validation, S.B., X.F., and J.F. wrote the manuscript with the help of S.M. for Methods.

## Data and code availability

The refined maps are deposited on EMDB and corresponding atomic models on PDB and will be publicly available as of the date of publication.

EMD-29681 (control apo-70S at 900ms), 29688 (i70S_HflX_-I), 29687 (i70S_HflX_-II), 29689 (i70S_HflX_-III), 29833 (consensus i70S_HflX_-I, 30S focused), 29834 (i70S_HflX_-I, 50S focused and 30S subtracted), 29724 (consensus i70S_HflX_-II, 30S focused), 29844 (i70S_HflX_-II, 50S focused and 30S subtracted), 29723 (consensus i70S_HflX_-III, 30S focused), 29842 (i70S_HflX_-III, 50S focused and 30S subtracted).

PDB-8G2U (control apo-70S at 900ms), 8G34 (i70S_HflX_-I), 8G31 (i70S_HflX_-II), 8G38 (i70S_HflX_-III).

