## Supplementary material for "Time resolution in cryo-EM using a novel PDMS-based microfluidic chip assembly and its application to the study of HflX-mediated ribosome recycling": Methods and SI

[3]Current address: Thermo Fisher Scientific, Oregon, USA

[#]These authors contributed equally

*Corresponding authors

†Lead Contact

### METHODS

#### Protein purification.

The pET28a-hflX plasmid (gift from Professor Ning Gao, Peking University, China) was transformed into *E. coli* BL21 Star™ (DE3)pLysS cells (Invitrogen) for HflX expression. Cells were incubated at 37°C until OD600 reached 0.6 and then induced with 0.6 mM isopropyl-β-D-thiogalactopyranoside (IPTG) (Sigma-Aldrich) for 12 hr at 16 °C. Cells were harvested and lysed in buffer 20 mM Tris-HCl, pH 7.6, 500 mM NaCl, 10 mM imidazole, 5 mM β-mercaptoethanol (βME), and 1 mM PMSF by ultrasonication. After centrifugation at 12,000 r.p.m. of cell lysates, supernatants were loaded onto a HisTrap HP Ni-NTA column (GE Healthcare) and eluted with buffer 20 mM Tris-HCl, pH 8.0, 500 mM NaCl, and 250 mM imidazole. HflX-containing fractions were pooled and concentrated using Amicon™ Ultra-4 Centrifugal Filter Units (MWCO 10kDa, MilliporeSigma™). TEV enzyme digestion was done to remove the histidine tag. Untagged HflX was further purified, and buffer exchanged using a gel-filtration column of Superdex 75 10/300GL (GE Healthcare) pre-equilibrated with buffer 20 mM Tris-HCl, pH 7.6, 500 mM NaCl, and 5 mM βME.

#### Isolation and purification of *E. coli* 70S ribosomes.

Log-phase *E. coli* MRE600 cells were harvested and washed with 20 mM Tris-HCl, pH 7.5, 10 mM $Mg(OAc)_2$, 100 mM $NH_4Cl$, and 5 mM βME buffer. Re-suspended cells in the same buffer were lysed with ultrasonication and centrifuged at 12,000 r.p.m. to remove cell debris. The supernatant containing crude ribosome was placed on the top of a 30% sucrose cushion (in 20 mM Tris-HCl, pH 7.5, 10 mM $Mg(OAc)_2$, 30 mM $NH_4Cl$, 5 mM βME) and pelleted down using Beckman 50.2 Ti rotor at 100,000 × g for 16 hr at 4°C. Clear ribosome pellet was re-suspended in 20 mM Tris-HCl, pH 7.5, 10 mM $Mg(OAc)_2$, 30 mM $NH_4Cl$, and 5 mM βME and homogenized for 1 hr in presence of 1 M $NH_4Cl$, after which the ribosomal preparation was centrifuged multiple times at 16,000 g and at 4 ºC for 20 min to remove all the non-specific proteins associated with ribosome. The supernatant containing ribosome was placed on the top of a 30% sucrose cushion (in 20 mM Tris-HCl, pH 7.5, 10 mM $Mg(OAc)_2$, 30 mM $NH_4Cl$, 5 mM βME) and pelleted down using Beckman 50.2 Ti rotor at 100,000 × g for 16 hr at 4°C. The pellet was dissolved in 20 mM Tris-HCl, pH 7.5, 10 mM $Mg(OAc)_2$, 30 mM $NH_4Cl$, and 5 mM βME and loaded on top of a 10 %–40 % linear sucrose gradient in 20 mM Tris-HCl, pH 7.5, 10 mM $Mg(OAc)_2$, 30 mM $NH_4Cl$, 5 mM βME buffer and centrifuged using Beckman swing-out SW 28 Ti rotor 28,000 r.p.m., 4 ºC, 1 hr 45 min. Fractions were collected using Gilson FC203B automated fraction collector. The 70S containing fractions were collected and sucrose was removed by exchanging with 20 mM Tris-HCl, pH 7.5, 10 mM $Mg(OAc)_2$, 30 mM $NH_4Cl$, and 5 mM βME buffer and concentrated to the desired amount using Amicon™ Ultra-4 Centrifugal Filter Units (MWCO 100 kDa, MilliporeSigma™). See Figure S1A.

#### Splitting assay.

1 µM 70S subunits were mixed with 1 µM HflX and 1mM GTP in 20 mM Tris-HCl, pH 7.5, 100 mM $NH_4Cl$, 10 mM $Mg(OAc)_2$, and 4 mM βME mixing buffer, and the reaction mixture was incubated at room temperature for 45 min. The mixtures were loaded onto a 10-40% (w/v) sucrose

cushion (prepared in mixing buffer) and centrifuged at 35,000 r.p.m. for 1 hr 15 min at 4 °C with an MLA-130 rotor (Beckman Coulter, Brea, California, USA). As a control, identical procedures were performed with the 70S and the same volume of mixing buffer; see Figure S1B.

**Co-sedimentation assay.**

1 μM 70S subunits were mixed with 1 μM HflX and 1mM GTP in 20 mM Tris-HCl, pH 7.5, 100 mM $NH_4Cl$, 10 mM $Mg(OAc)_2$, and 4 mM βME mixing buffer, and the reaction mixtures were incubated at room temperature for 5 min. The mixtures were loaded onto a 30% (w/v) sucrose cushion (prepared in mixing buffer) and centrifuged at 85,000 r.p.m. for 1 hr at 4 °C with an MLA-130 rotor (Beckman Coulter). As a control, identical procedures were performed with 70S and same volume of mixing buffer. Supernatants were carefully removed, and pellets were resuspended in mixing buffer and analyzed by 12% SDS-PAGE. See Figure S1C.

**The apparatus for Time-resolved (TR) cryo-EM sample preparation.**

The apparatus for accommodating and controlling the microfluidic chip (Figure S2) was built by Dr. Howard White[1,2], but some new features were added as follows: an environmental chamber for maintaining the temperature and humidity[3], and a novel microfluidic chip assembly for depositing time-resolved reaction product on the EM-grid. In addition, we now use two separate gas pumping systems to control the plunger and microsprayer, thereby producing more stable plunging motion and solution atomization, respectively, compared to the original apparatus with a single gas pumping system.

The chip is mounted in the environmental chamber, with the nozzle of the microsprayer facing the cryo-EM grid. Two solutions of interest are fed into the micromixer of the microfluidic chip through 1/16-inch polyetheretherketone (PEEK) capillary tubing (125 μm inner diameter) and 2.0-μm PEEK microfilters, controlled by a computer-assisted liquid-pumping and grid-plunging apparatus designed by H. White and co-workers[1,2]. Each EM grid to be tested is mounted on sharp-tip tweezers. The other end of the tweezers is mounted on a pneumatic motor, which is controlled by computer. For detailed information on the plunging machine, see the work by White et al.[2]. As cryogen, liquid ethane is used. The averaged plunging speed can be controlled by the gas pressure, and 40 psi was used in this study to generate a plunging velocity of 1.9 m/s. The distance from the sprayer nozzle to the surface of the liquid ethane surface can be adjusted down to a minimum value of 10 mm. The horizontal distance sprayer-grid was fixed at 3.5 mm. The ambient conditions in the environmental chamber can be maintained in the ranges of 22°C–25°C temperature and 85%–95% relative humidity. Compared with our previous design[3], the newly-designed micro-sprayer works better even at the lower pressure of 8 psi without producing the previously observed dripping problem[3]. In this work, we used 8 psi as the working gas pressure throughout, which ensures stable, reproducible spray.

**High-efficiency SAR PDMS-based micromixer.**

To efficiently and fast mix the solutions of interest, we used the 3D SAR PDMS-based micro-mixer[4] (or 3D crossing micro-mixer), which is superior to the planar micro-mixer (for example, the planar butterfly-type micromixer used in Lu et al.'s design[5] adopted in our previous work[6-10]).

In order to save computing time in the simulation, we simulated the mixing performance using the first five mixing units only and obtained the result summarized in Figure S3. As shown there, when the total flowrate ranges between 1 to 6 μL/s, the mixing efficiency, after passing five mixing units, is always higher than 90%. This working flow-rate range cannot be achieved with the butterfly-type micro-mixer of Lu, et al.'s design[5], whose performance at low flowrate is limited.

To measure the mixing efficiency of this micro-mixer, we conducted experiments by injecting DI water and fluorescent water into its two separate inlets. The mixing efficiency $E$ can be quantitatively evaluated[11] as $E = (1 - \sqrt{1/n \sum (I_i - I_{av})^2} / I_{av}) \times 100\%$, where $I_i$ is the fluorescent intensity in pixel $i$, $n$ is the total number of pixels, and $I_{av}$ is the average intensity of $n$ pixels. For unmixed fluids, $E = 0$, and for completely mixed fluids, $E = 1$. Usually, $E > 90\%$ is taken to indicate excellent mixing performance.

The results of our mixing experiments are shown in Figure 1D. We observed that at a total flow rate of 1 μL/s, the fluorescence intensity is not perfectly distributed at the mixer outlet, indicating that the mixing was inefficient under this condition. When the flowrate was increased to 6 μL/s, more striations appeared at the mixer outlet because of the strong rotation and splitting of the contact surface between two fluids, indicating that the mixing efficiency was greatly enhanced. Based on the good agreement between experimental and simulation results, we find that in the whole range of total flowrates from 2 to 6 μL/s, the mixing efficiency at the outlet exceeds 90%. From this range, 6 μL/s was chosen as the working flowrate for this study to achieve both the shortest mixing time and efficient mixing performance.

**$SiO_2$-coating inside the micromixer for mitigating protein adsorption.**

$SiO_2$ coating was performed inside the PDMS micro-mixer channel to prevent proteins of the sample from sticking to the PDMS plastic. The measurements show that up to 50% of the protein is lost without coating. Adverse effects of this problem include uncontrolled changes in concentration of the two mixing components during the use of the micromixer, which alter the stoichiometry of the reaction, rendering the results of time-resolved study invalid.

For $SiO_2$ coating, plasma-enhanced chemical vapor deposition (PECVD) was performed to deposit a thin $SiO_2$ layer onto the interior PDMS micro-mixer channel walls using the PlasmaPro®NGP80 system (Oxford Instruments, Abingdon, UK). High radio frequency (RF) power (50W) was used to create plasma inside the process chamber. During plasma treatment, the source gases used for plasma were $N_2O$ (710 sccm) and $SiH_4$ (170 sccm), the vacuum pressure was 200 mTorr, the substrate temperature was 300 °C, and the coating strike lasted for 20 min. The $SiO_2$ layer is around 1.9 μm in thickness, as measured on Filmetrics F20. The reaction formula is as below for $SiO_2$ layer formation under plasma conditions:

$$2N_2O + SiH_4 = SiO_2 + 2N_2 + 2H_2 \qquad (1)$$

Protein adsorption assessment with *E. coli* 70S ribosome was performed to compare the properties of differently treated chip assemblies: without coating, coated with n-dodecyl-β-D-maltoside (DDM) coating, or coated with $SiO_2$. DDM is an alkyl polyglucoside, a very mild non-ionic surfactant which can be used for improving the hydrophilicity of the surface of PDMS. For each of these chip assemblies an experiment was performed by passing through it a sample of 70S ribosomes at least six times, then spraying it out and collecting it for concentration measurement by spectrophotometric analysis using NanoDrop® (Thermo Fisher Scientific, Waltham, MA, USA). The absorbance values at 260 nm ($A_{260}$) for the initial concentration as control was 0.966 on average (where $A_{260}$ = 1 = 20nM). The $A_{260}$ values for the samples using chips with different coating methods are shown in Table S2. The measurements of ribosome concentration after passing the sample through the chips show (Figure 1H) that 94% of the initial concentration is retained using the $SiO_2$-coated chip, while only 54% and 60% of the initial concentration are retained without coating or with DDM coating, respectively, demonstrating that the $SiO_2$ coating can effectively mitigate the problem of protein adsorption.

**Choice of microcapillary tubing as the reactor for stable reaction time control.**

In this study, we are using micro-capillary tubing with circular transverse section as micro-reactor. In principle, the fluid flow is subject to the no-slip boundary condition at the walls. The hydrodynamic entrance length is 0.05*Re*D, where *Re* is the Reynolds number and D is the diameter of the tubing. After passing the micro-capillary over a distance equal to the hydrodynamic entrance length, the velocity profile of the fluid flow is fully developed into a parabolic profile. So, a fully developed velocity profile (FDVP) for a circular tubing can be expressed in the following equation[12],

$$V(y,z) = 2\bar{V}\left(1 - \frac{(y-A)^2 + (z-B)^2}{r^2}\right) \tag{2}$$

where $\bar{V}$ is the mean velocity, (*A, B*) are the coordinates of the center of the inlet, and *r* is the inner radius of micro-reactor. The mean velocity is determined by the volume of the tubing and the volumetric flowrate. For a tubing with $r$ = 75 µm, the hydrodynamic entrance length is 0.38 mm, which means that the fluid flow can be easily developed to be stable after 1 ms based on the parabolic profile (Figure S4)

**Redesign of the micro-sprayer.**

In our previous work[3], the micro-sprayer (left in Figure S5) was designed and fabricated to generate a three-dimensional cone plume of sprayed droplets. This micro-sprayer contains mainly an inner tubing serving as a liquid injector and an outer tubing as gas nozzle, which are both accommodated in the PDMS slab. Orifices of inner and outer tubing were not aligned in that design. Subsequently, in practical experiments, we found out that part of the solution was dripping from the orifice when lower gas pressure was used. To solve this dripping problem, we aligned the orifices of the inner and outer tubings on the same plane and also centered them precisely, as shown in Figure 5S. After this redesign, the micro-sprayer generated a cone of droplets at gas pressure of 8 psi and flowrate of 6 µL/s without exhibiting the dripping problem. With this micro-sprayer, the spray is found to be stable and reproducible, at pressure conditions low enough to prevent damage to the EM grid.

**Assembly and disassembly of the TR chip.**

For the fabrication of the entire chip assembly, we developed a modular strategy such that it can be customized for individual experiments by assembling it from three modules (mixer, reaction channel, sprayer), allowing each of its parts to be reused after disassembly (Figure S6). The advantage of the modular design is that any of the three elements of the chip that is found not functioning can be readily replaced by a new one. This is also convenient for cleaning the microreactor channel through oxygen plasma treatment before each new reaction experiment. Normally, the microfluidic chip assembly is used for a duration of one month.

**Estimation of reaction time achieved using this TR method.**

As in our previous mixing/plunging method[5], the total reaction time consists of the mixing time in the micro-mixer, the reaction time in the micro-capillary reactor, the spraying time, the plunging time, and the reaction time during the vitrification process. For our current method, each time is separately calculated below.

1. The mixing time in the micromixer, $\mathrm{t_m}$. The mixing time is estimated by the equation[13]: $\mathrm{t_m} = V_m/U_f$, $V_m$ is the volume of micromixer, $U_f$ is the total flowrate of the solutions introduced into the micromixer. As measured in the 3D modelling software FreeCAD, $V_m$ is 280 nL, and $U_f$ is 6 µL/s, so the mixing time is estimated to be ~0.47 ms.
2. The reaction time in the microcapillary reactor, $\mathrm{t_r}$. Because of the parabolic profile of velocity distribution in the capillary tubing, the real residence time for the reaction is in a time range. To simplify, we estimate the reaction time in the tubing based on the mean velocity and the volume of the tubing: $\mathrm{t_r} = L_r/\bar{V}$ , where $L_r$ is the length of the microcapillary reactor, $\bar{V}$ is the mean velocity of the fluid, which is equivalent to $U_f/S$, and $S$ is the transversal area of the micro-capillary.
3. The flying time of the droplets from the micro-sprayer orifice to the EM grid, $\mathrm{t_f}$. The Sauter mean diameter (SMD) as the average of the droplets can be estimated by the following expression[14,15],
$$SMD = 0.95\left[\frac{(\sigma_l \dot{m}_l)^{0.33}}{V_r \rho_l^{0.37} \rho_g^{0.30}}\right]\left[1+\frac{\dot{m}_l}{\dot{m}_g}\right]^{1.70} + 0.13\mu_l\left[\frac{D}{\sigma_l \rho_l}\right]^{0.5}\left[1+\frac{\dot{m}_l}{\dot{m}_g}\right]^{1.70} \quad (3)$$
where $\dot{m}$ is the mass flow rate, and subscripts $g$ and $l$ denote gas and liquid. Suppose that water is used for atomization, for which viscosity $\mu_l$, surface tension $\sigma_l$, density $\rho_l$ are $0.89\times10^{-3}$ Pa·s, 0.072 N/m, $1\times10^3$ kg/m$^3$, respectively. $D$ is diameter of the inner tubing of the micro-sprayer, which is 75 µm. For our study, the flow rate of the liquid is 6 µL/s and the mass flow rate $\dot{m}_l$ of liquid is 0.0216 kg/h. The liquid velocity at the micro-sprayer orifice[16] is $16\dot{m}_l/5\rho_l\pi D^2$, which is 1.1 m/s. At 8 psi, the volumetric flow rate measured is ~0.2 L/min, and the density $\rho_g$ of $N_2$ is 1.71 kg/m$^3$ at room temperature, so the mass flow rate $\dot{m}_g$ of $N_2$ gas is 0.02 kg/h and the gas velocity *is* $V_g$= ~31.6 m/s. $V_r$ is the relative velocity between the gas and liquid, $V_r = V_g - V_l$; for estimation, we use the liquid velocity at the microsprayer orifice as $V_l$. We also have the Weber number based on the SMD,
$$We_{SMD} = \frac{\rho_l V_l^2 SMD}{\sigma_l} \quad (4)$$
The velocity of the droplet depends on the ratio of dynamic force to surface tension force, and it is governed by the following relation[17],
$$We_c = \frac{\rho_g (V_g - V_d)^2 SMD}{\sigma_l} \approx We_{SMD} \quad (5)$$
Then the averaged velocity of the droplets $V_d$ can be estimated, which is around 6.4 m/s, so that we can estimate the mean flying time of the droplets to be $\mathrm{t_f} = D_f/V_d$, where $D_f$ is the distance from the micro-sprayer orifice to the EM grid, fixed at 3.5 mm for our implementation, and thus we obtain a mean droplet flying time of ~0.55 ms.

4. The plunging time, $t_p$. The plunger is pneumatic, and the plunging speed $V_p$ =1.9 m/s at $N_2$ gas pressure of 40 psi. So, the plunging time $t_p = H_p / V_p$, where $H_p$ is the plunging height from the micro-sprayer orifice to the liquid ethane surface.
5. The reaction time during the vitrification process, $t_v$. We assume that at 0°C (273.15K) the reaction is slow, and the reaction time can be neglected when the temperature is going down to below 0°C during the vitrification process. And we have the critical cooling rate (CCR)[18] which is ~$10^5$ K/s. So, we can estimate the reaction time in the vitrification process is $t_v = (T_{room} - 273.15)/\mathrm{CCR}$, which amounts to ~0.23 ms.

Thus in total, the reaction time can be estimated as $t = t_m + t_r + t_f + t_p + t_v$ . In practice, the shortest reaction time we are able to achieve is ~10 ms, when the microreactor tubing is 5 mm in length and 75 μm in diameter, and the plunging height is 10 mm. By varying the micro-reactor tubing length, the reaction time can be tuned from 10 to 1000 ms. For the application in the HflX study we implemented only tubings for four reaction time points, and all the parameters about the chip assemblies are listed in Table S1. Based on the above-mentioned time estimates, the reaction times can be achieved at 10, 25, 141, and 899 ms, respectively (For simplification, we use rounded figures of 140 and 900 ms in the following).

**Preparation of the time-resolved cryo-grids using the TR chip.**

Quantifoil Cu R0.6/1 grids with 300 mesh size were subjected to glow discharge with air for 30 s using a PELCO easiGlow cleaning system set to a plasma current of 15 mA, to make the carbon film surface negatively charged (hydrophilic), which allows aqueous solutions to spread easily. During the experiment, the microfluidic chip was mounted in an environmental chamber[6], in which the temperature and humidity were kept at 22 °C ~24 °C and 90%~95%, respectively. For each of the four time points (10 ms, 25 ms, 140 ms, 900 ms) and the control experiment, the 70S, HflX, and GTP were diluted to 1 μM, 5 μM, and 1 mM, respectively, with 20 mM Tris-HCl, pH 7.5, 100 mM $NH_4Cl$, 10 mM $Mg(OAc)_2$, and 4 mM βME mixing buffer. Solution A: 1 μM 70S in mixing buffer, and Solution B: 5 μM of HflX with 1 mM GTP in the same buffer were introduced into the different microfluidic chips at a flow rate of 3 μL/s for each, such that they were mixed efficiently and sprayed onto a plasma-treated grid. The resulting concentrations of the 70S ribosome and HflX after effective mixing in our microfluidic chip were around 0.5 μM and 2.5 μM, respectively (Concentrations were measured separately for each component after collection from the chip.) After the reaction product was sprayed onto the grid, the latter was immediately plunged into liquid ethane for the vitrification of the sample. Based on our previous study[3], the gas pressure for atomization was controlled at 8 psi to generate properly sized droplets for data collection. Each grid was stored in liquid nitrogen dewar until it was ready for imaging.

**Preparation of EM grids for control experiment for Apo-70S using the TR chip.**

Solution A: 1 μM 70S in mixing buffer and Solution B: only the mixing buffer, were introduced into the 900 ms microfluidic chip. The next steps were the same as described above.

**TR cryo-EM data collection.**

All the data from both the TR and control experiments were collected using a 300 kV Titan Krios (Thermo Fisher Scientific, Waltham, MA) equipped with a K3 direct detector camera (Gatan, Pleasanton, CA). For the blotted grids from the control experiment, the data collection followed our previous procedure[6-8,10], and was done automatically using the Leginon software[19]. For the TR-grids, on the square targeting, the positions of collectible droplets need to be picked up from the atlas based on our previous work[7,10], and on the hole targeting, the thick-ice area needs to be avoided based on the intensity on the hole images as shown in Figure S7, so these two steps can be finished manually. Focusing was performed on the carbon foil before each exposure. After focusing twice, an exposure was immediately taken with an image shift producing a beam-tilt smaller than 0.005 mrad. For TR grids, in the exposure mode, the movie stacks were recorded within a defocus range of 1.0 to 2.5 µm on a Gatan K3 Summit direct detector camera combined with the Gatan Bioquantum energy filter (slit width of 20 eV), operating in counting mode with an effective magnification of 105,000×, equivalent to 0.83 Å per pixel. Images were composed of 50 frames that were exposed for a total of 2.5 s, corresponding to a total dose of 58 $e^{-}/Å^{2}$. Some representative micrographs from TR grids are shown in Figure S9.

**Cryo-EM data processing.**

A flow-chart of the data processing containing the steps is shown in Figure S10. 3452, 3598, 3530, and 3603 good micrographs were selected from TR experiments at 10, 25, 140, and 900 ms, respectively, for further data processing. The beam-induced motion of the sample was corrected using the MotionCor2 program[20]. The contrast transfer function (CTF) of each micrograph was estimated using the CTFFIND4[21]. Particle picking was performed using Topaz[22]. Good particles were selected by 2D classification and trained using 20,000 particles. Autopicking using the trained topaz model yielded 1,001,596 particles from all combined good micrographs (total: 14,183). Particles picked by Topaz were subjected to 2D classification for a further selection of good particles, which yielded 802,562 particles. All particles were pooled together and used for 3D initial model generation followed by 3D auto-refinement, applying C1 symmetry in Relion 4[23]. CTF refinements were done to correct for magnification anisotropy, fourth-order aberrations, per-particle defocus, and per-particle astigmatism, followed by another 3D auto-refinement. Then 3D classification was performed on the entire pooled dataset[7,10] without alignment, using the angular information from the previous refinement step. The 3D classification produced seven distinct classes, which we named (1) rotated 70S without HflX ($r70S_{noHflX}$, 56,894 particles); (2) non-rotated 70S without HflX ($nr70S_{noHflX}$, 96,898 particles); (3) 70S like intermediate-I with HflX ($i70S_{HflX}$-I, 140,682 particles); (4) 70S like intermediate-II with HflX ($i70S_{HflX}$-II, 138,296 particles); (5) 70S like intermediate-III with HflX ($i70S_{HflX}$-III, 113,038 particles); (6) 50S with HflX ($50S_{HflX}$, 62,558 particles); and (7) 30S (58,952 particles) with total 667,318 particles. The reproducibility of the classes was checked using the previously reported method[10]. Percentages of particles falling in each class with respect to the total of 667,318 are shown in Figure 3M. Particles from the $50S_{HflX}$ class were further subjected to 3D auto-refinement, and the post-processing was done with RELION4 and DeepEMhancer[24]. The angular distributions for the reconstructions are shown in Figures S13G-J. For the intermediate states, from the 3D classes (Figure S10) the 30S portion was first separated by masking and focused refinement was performed, then the projected density corresponding to the 30S subunit was subtracted from the refined particles, and focused refinement was done on the 50S subunit part. All the FSC resolutions estimations are contained in Figure S12. For the refinement of the HflX density region, particles from each intermediate state and the $50S_{HflX}$ subunit were individually masked and 3D classification was performed on the masked-out HflX density without alignment. Further, Ewald sphere correction[25] was performed on the two resulting half-maps using relion_reconstruct. In the control TR experiment for Apo-

70S, the same above-mentioned processing steps were followed with 140,338 particles. The corresponding particle numbers for non-rotated and rotated apo-70S are 75,552 and 64,786, respectively. Here we found only the 70S state, so for the purpose of the particle percentage calculation, it was considered as a 100% population. To build models for the 70S ribosome class reconstructions and HflX, the pdb id:6XE0[26] for 30S, pdb id:6XZ7[27] for 50S, pdb id: 4PYG[28] for GTP, pdb id: 1GIT[29] for GDP-Pi, and pdb id: 5ADY[30] for HflX were selected as starting models. Models were first refined using Phenix real-space refinement[31]. Residues that did not fit correctly into the map were manually placed using COOT[32]. Further model validations were done using Phenix Comprehensive Validation (cryo-EM) and tabulated in Table S3.

**Determining the position vector of the unique point through which the rotation axis passes**

The tool we previously developed[33] for determining the rotation axis, using the absolute orientation method produces a closed-form solution of the coordinate axis transformation in the form of a unit quaternion. A description of the algorithm and details of implementation are provided in our previous work[33]. The unit quaternion encodes the rotation axis and rotation angle, which can be obtained using the angle-axis form of the unit quaternion. However, although the translation from the initial to the transformed centroids can be computed easily, the unit quaternion representing the rotation does not contain the position vector of the unique point $\vec{C}_\perp$ (Figure 6L) through which the rotation axis $\hat{l}$ passes. For determining $\vec{C}_\perp$, we use the theory of screw axis of spatial displacement[34-37] which has also been used in a few other applications in the literature[38-40]. Here we provide a formulation of the rigid-body motion along with the derivation of $\vec{C}_\perp$ (Figure 6L) below.

We use the position vectors of the start and the end position of the rigid body center of mass as depicted in Figure 6L. Let $\vec{t}_\parallel$ and $\vec{t}_\perp$ be the components of the translation $\vec{t}$, that are parallel and perpendicular respectively to the rotation axis $\hat{l}$. Then we have,

$$\vec{t}_\parallel = (\hat{l} \cdot \vec{t})\, \hat{l} \tag{6}$$

$$\Rightarrow \vec{t}_\perp = \vec{t} - \vec{t}_\parallel = \vec{t} - (\hat{l} \cdot \vec{t})\, \hat{l} \tag{7}$$

Now,

$$\hat{y}_b = \frac{\vec{t}_\perp \times \hat{l}}{\|\vec{t}_\perp \times \hat{l}\|} = \frac{\vec{t}_\perp \times \hat{l}}{\|\vec{t}_\perp\|\, \|\hat{l}\| \left|sin\left(\frac{\pi}{2}\right)\right|} = \frac{\vec{t}_\perp \times \hat{l}}{\|\vec{t}_\perp\| \cdot 1 \cdot 1} = \frac{\vec{t}_\perp \times \hat{l}}{\|\vec{t}_\perp\|} \tag{8}$$

Also from Figure 6,

$$\frac{\frac{1}{2}\, \|\vec{t}_\perp\|}{\|\vec{y}_b\|} = \tan\frac{\theta}{2}$$

$$\Rightarrow \|\vec{y}_b\| = \frac{1}{2}\cot\frac{\theta}{2}\, \|\vec{t}_\perp\| \tag{9}$$

Then we have from equations (8) and (9),

$$\vec{y}_b = \|\vec{y}_b\|\, \hat{y}_b = \frac{1}{2}\cot\frac{\theta}{2}\, \|\vec{t}_\perp\| \frac{\vec{t}_\perp \times \hat{l}}{\|\vec{t}_\perp\|}$$

$$\Rightarrow \vec{y}_b = \|\vec{y}_b\|\, \hat{y}_b = \frac{1}{2}\cot\frac{\theta}{2}\; \vec{t}_\perp \times \hat{l} \tag{10}$$

Now,

$$\overrightarrow{CA_p} + \frac{1}{2}\vec{t}_\perp = \vec{y}_b$$

$$\Rightarrow \overrightarrow{CA_p} = \vec{y}_b - \frac{1}{2}\vec{t}_\perp \tag{11}$$

Next,

$$\overrightarrow{OA_p} = \overrightarrow{OC} + \overrightarrow{CA_p}$$

$$\Rightarrow \overrightarrow{OC} = \overrightarrow{OA_p} - \overrightarrow{CA_p}$$

Then using equation (11),

$$\overrightarrow{OC} = \overrightarrow{OA_p} - \left(\vec{y}_b - \frac{1}{2}\vec{t}_\perp\right) \tag{12}$$

Now,

$$\overrightarrow{OA_p} = \overrightarrow{OA} - \overrightarrow{A_pA} \tag{13}$$

Also, let the angle between $\overrightarrow{OA}$ and the plane be $\alpha$, so the angle between $\overrightarrow{OA}$ and $\hat{l}$ is $\frac{\pi}{2} - \alpha$, then we have:

$$\overrightarrow{OA} \cdot \hat{l} = \|\overrightarrow{OA}\|\|\hat{l}\| \cos\left(\frac{\pi}{2} - \alpha\right)$$

$$= \|\overrightarrow{OA}\| \cdot 1 \cdot \sin\alpha = \|\overrightarrow{OA}\| \sin\alpha$$

Also, we have $\|\overrightarrow{OA}\| \sin\alpha = \|\overrightarrow{A_pA}\|$

$$\Rightarrow \overrightarrow{OA} \cdot \hat{l} = \|\overrightarrow{A_pA}\| \tag{14}$$

Now, the vector,

$$\overrightarrow{A_pA} = \|\overrightarrow{A_pA}\|\,\hat{l}$$

Using equation (14)

$$\overrightarrow{A_pA} = \left(\overrightarrow{OA} \cdot \hat{l}\right)\,\hat{l} \tag{15}$$

Using equations (13) and (15), we have

$$\overrightarrow{OA_p} = \overrightarrow{OA} - \overrightarrow{A_pA} = \overrightarrow{OA} - \left(\overrightarrow{OA} \cdot \hat{l}\right)\,\hat{l} \tag{16}$$

Therefore, using equations (10), (12) and (16), we have

$$\overrightarrow{OC} = \overrightarrow{OA_p} - \left(\vec{y}_b - \frac{1}{2}\vec{t}_\perp\right) = \overrightarrow{OA} - \left(\overrightarrow{OA} \cdot \hat{l}\right)\,\hat{l} - \left(\frac{1}{2}\cot\frac{\theta}{2}\;\vec{t}_\perp \times \hat{l} - \frac{1}{2}\vec{t}_\perp\right)$$

Now denoting $\overrightarrow{OC}$ by $\vec{C}_\perp$ , $\overrightarrow{OA}$ by $\vec{x}_0$ , $\overrightarrow{OB}$ by $\vec{x}_1$ we have,

$$\vec{C}_{\perp} = \vec{x}_0 - (\vec{x}_0 \cdot \hat{l})\ \hat{l} + \frac{1}{2}\left[\vec{t}_{\perp} - \cot\frac{\theta}{2}\ \ \vec{t}_{\perp} \times \hat{l}\right]$$

$$\Rightarrow \vec{C}_{\perp} = \vec{x}_0 - (\vec{x}_0 \cdot \hat{l})\ \hat{l} + \frac{1}{2}\left[\vec{t}_{\perp} + \cot\frac{\theta}{2}\ \ \hat{l} \times \vec{t}_{\perp}\right] \tag{17}$$

Now from equation (2), we have

$$\vec{t}_{\perp} = \vec{t} - (\hat{l}\cdot\vec{t})\,\hat{l} = \hat{l} \times (\vec{t} \times \hat{l})$$

and

$$\hat{l} \times \vec{t}_{\perp} = \hat{l} \times \left(\vec{t} - (\hat{l}\cdot\vec{t})\,\hat{l}\right) = \hat{l} \times \vec{t} - (\hat{l}\cdot\vec{t})\,\hat{l} \times \hat{l} = \hat{l} \times \vec{t} \qquad [as\ \hat{l} \times \hat{l} = 0]$$

Letting $\vec{b} = \tan\frac{\theta}{2}\ \hat{l}$ we can rewrite,

$$\vec{C}_{\perp} = \vec{x}_0 - (\vec{x}_0 \cdot \hat{l})\ \hat{l} + \frac{\vec{b} \times \vec{t} - \vec{b} \times (\vec{b} \times \vec{t})}{2\vec{b}\cdot\vec{b}} \tag{17a}$$

Next also we have $\vec{t} = \vec{x}_1 - \vec{x}_0$

Therefore, equation (17) can be re-written in terms of the position vectors $\vec{x}_0$ & $\vec{x}_1$

$$\vec{C}_{\perp} = \vec{x}_0 - (\vec{x}_0 \cdot \hat{l})\ \hat{l} + \frac{1}{2}\left[\vec{t} - (\hat{l}\cdot\vec{t})\,\hat{l}\ + \cot\frac{\theta}{2}\ \ \hat{l} \times \vec{t}\right]$$

$$= \vec{x}_0 - (\vec{x}_0 \cdot \hat{l})\ \hat{l} + \frac{1}{2}\left[(\vec{x}_1 - \vec{x}_0) - \left(\hat{l}\cdot(\vec{x}_1 - \vec{x}_0)\right)\hat{l}\ + \cot\frac{\theta}{2}\ \ \hat{l} \times (\vec{x}_1 - \vec{x}_0)\right]$$

$$= \vec{x}_0 - (\vec{x}_0 \cdot \hat{l})\ \hat{l} + \frac{1}{2}\left[(\vec{x}_1 - \vec{x}_0) - \left(\hat{l}\cdot(\vec{x}_1 - \vec{x}_0)\right)\hat{l}\ + \cot\frac{\theta}{2}\ \ \hat{l} \times (\vec{x}_1 - \vec{x}_0)\right]$$

$$= \left[\vec{x}_0 + \frac{1}{2}(\vec{x}_1 - \vec{x}_0)\right] - \left[(\vec{x}_0 \cdot \hat{l})\ \hat{l} + \frac{1}{2}\left(\hat{l}\cdot(\vec{x}_1 - \vec{x}_0)\right)\hat{l}\right] + \frac{1}{2}\cot\frac{\theta}{2}\ \hat{l} \times (\vec{x}_1 - \vec{x}_0)$$

$$\Rightarrow \vec{C}_{\perp} = \frac{(\vec{x}_1 + \vec{x}_0)}{2} - \left(\hat{l}\cdot\frac{(\vec{x}_1 + \vec{x}_0)}{2}\right)\hat{l} + \frac{1}{2}\cot\frac{\theta}{2}\ \hat{l} \times (\vec{x}_1 - \vec{x}_0) \tag{18}$$

As we can see, the position vector $\vec{C}_{\perp}$ of the unique point on the rotation axis $\hat{l}$ can be expressed in terms of the initial and transformed position vectors $\vec{x}_0$ & $\vec{x}_1$, respectively, specifically in terms of the mid-point $(\vec{x}_0 + \vec{x}_1)/2$ and the translation vector $\vec{x}_1 - \vec{x}_0$. This provides a precise location estimate of the unique point through which the screw axis passes.

We can obtain (derivation not shown here) the exact same expression for $\vec{C}_{\perp}$ as in equation (18) using the Rodrigues displacement equation[41-44] for the rigid body motion as described in Figure 6L.

Note that the expression for $\vec{C}_{\perp}$ in equations (17) and (17a) contains the extra term $\vec{x}_0 - (\vec{x}_0 \cdot \hat{l})\ \hat{l} = \hat{l} \times (\vec{x}_0 \times \hat{l})$. This represents a generalized form, which allows the introduction of an arbitrary reference origin. The remaining terms in equations (17) and (17a) match with the previous derivation in the literature[45,46].

In this way we computed the rotation axis $\hat{l}$ and also the point with position vector $\vec{C}_{\perp}$ for the motion of the 30S subunit with respect to the large subunit of the ribosome (for Axis-I and -II). (Figure 6D-I). We have also computed the rotation axis for the 30S head rotation (Axis-III) with respect to the 30S subunit body (Figure S14C).

**Molecular Dynamics (MD) Simulation.**

MD simulation was performed on free HflX only, based on the method previously described[47,48]. The HflX model was taken from pdb id: 5ADY. The length of the simulation was 1000 ns. For the determination of flexibility of HflX domains, order parameters ($S^2$) were calculated as described previously[47,48]. Backbone N-H vectors were selected to calculate $S^2$ over the period of trajectory, which represents the dynamics of protein, with a value of 1 indicating complete rigidity and a value of 0 representing enhanced dynamics.

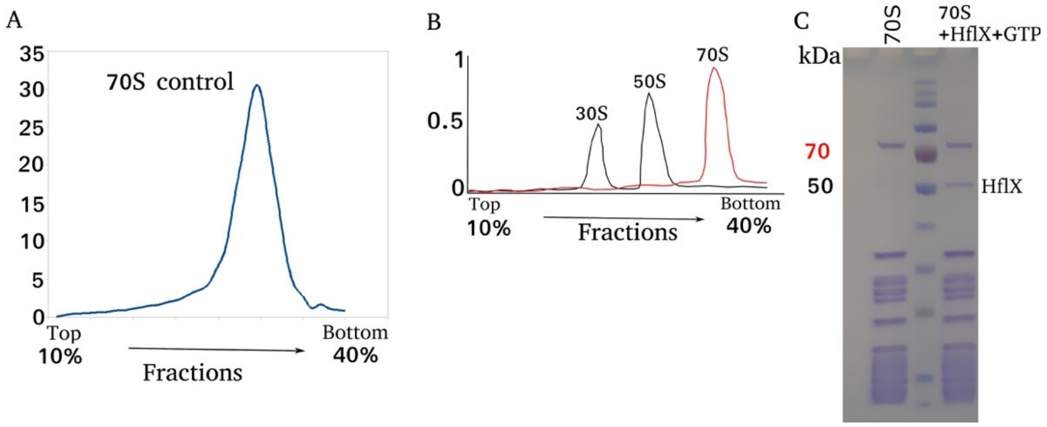


**Figure S1 | Activity check of the 70S and HflX.** (**A**) Sucrose gradient profile of the 70S purified from MRE600. (**B**) Sucrose gradient profile of dissociated 50S and 30S from 70S in presence of HflX and GTP, and shown in black line. Control 70S without HflX and GTP is shown in the red line. (**C**) SDS PAGE gel profile of control 70S (1st lane), 70S+HflX+GTP complex (3rd lane), and standard protein marker (2nd lane) representing the outcome of the co-sedimentation assay.

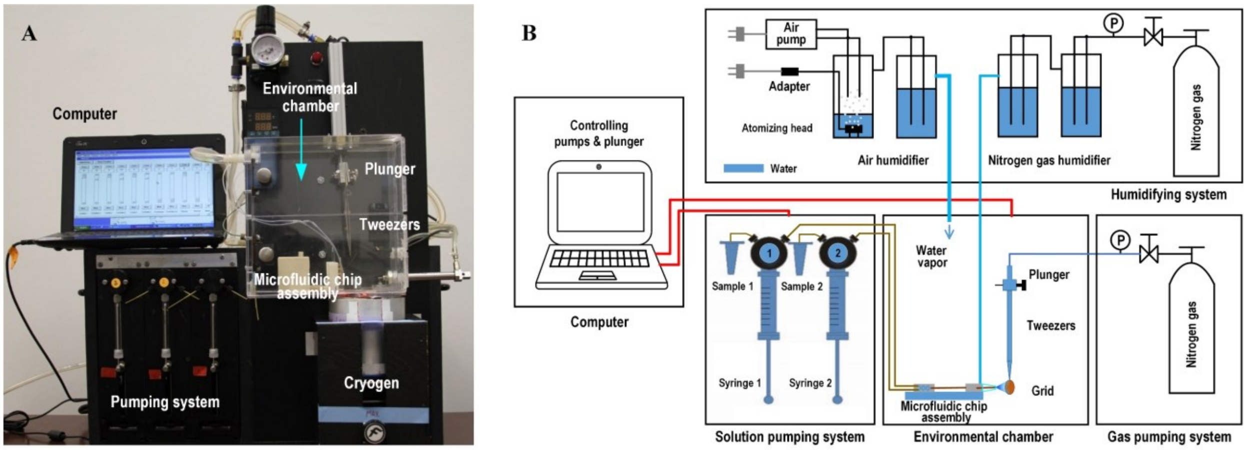


**Figure S2 | Time-resolved (TR) cryo-EM apparatus for sample preparation.** (**A**) Photograph of the setup, including computer, pumping system, pneumatic plunger, and environmental chamber. (**B**) Schematics of apparatus.

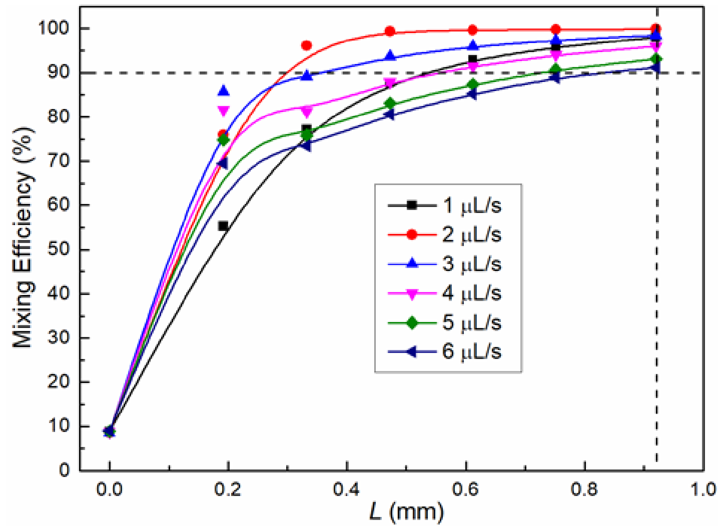


**Figure S3 | Result of mixing simulation for the 3D SAR PDMS-based micro-mixer**. Mixing efficiency along the outflow direction under different total flowrates, where L = 0.92 mm represents the position of the outlet.

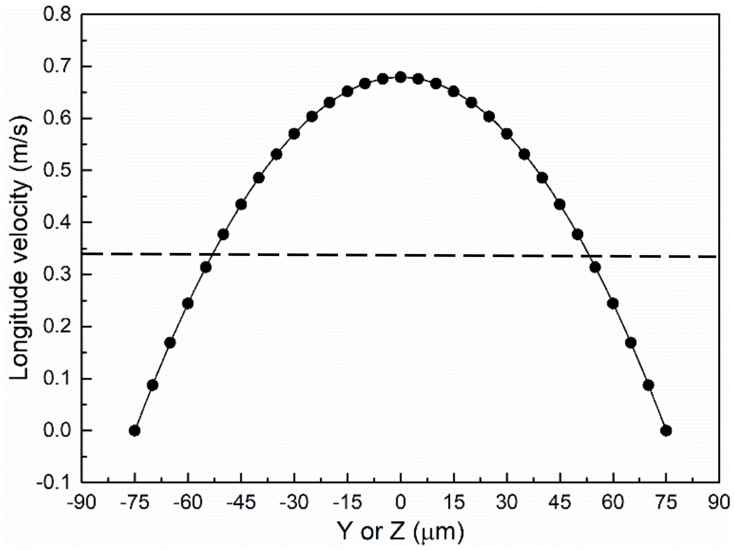


**Figure S4 | Parabolic profile in capillary tubing.** Numerically calculated velocity profile along the horizontal (y-axis) and vertical (z-axis) centerline of the inlet channel at different coordinates x for the fully developed case, with $\bar{V}$ = 0.34 m/s (*Re* = 51). The dashed line represents the mean velocity.

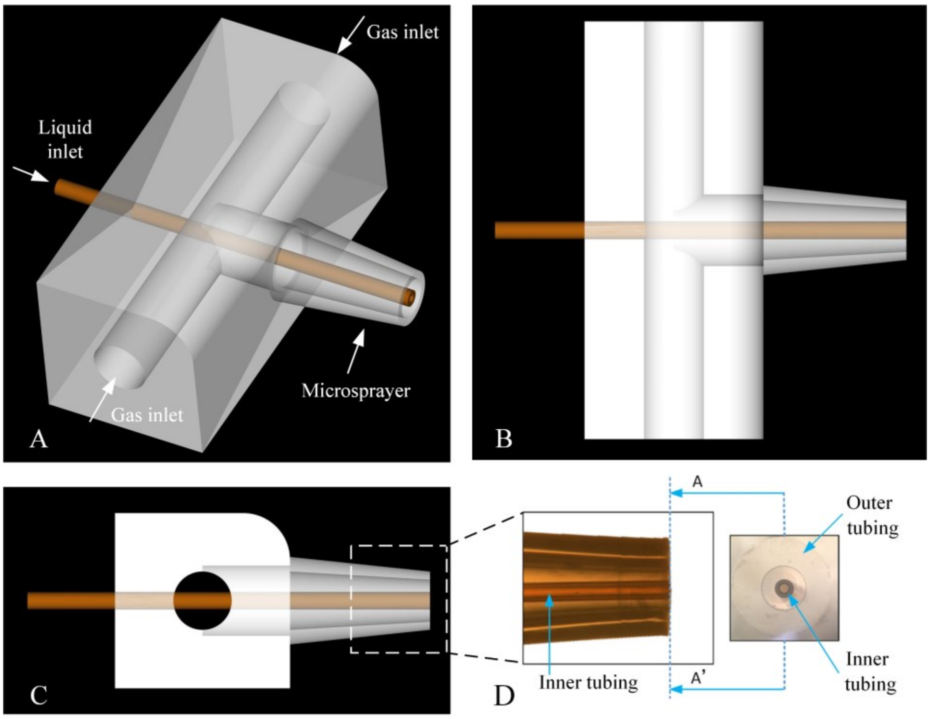


**Figure S5 | Design of the microsprayer.** (A), (B), and (C) 3D model of micro-sprayer in oblique, top, and side views. (D) Front portion of the fabricated micro-sprayer in side view.

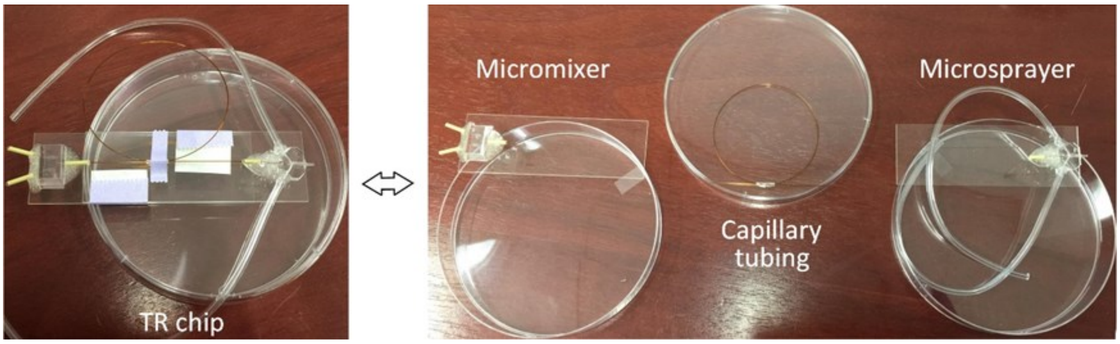


**Figure S6 | (Left) Complete TR chip assembly. (Right) Its three modules.**

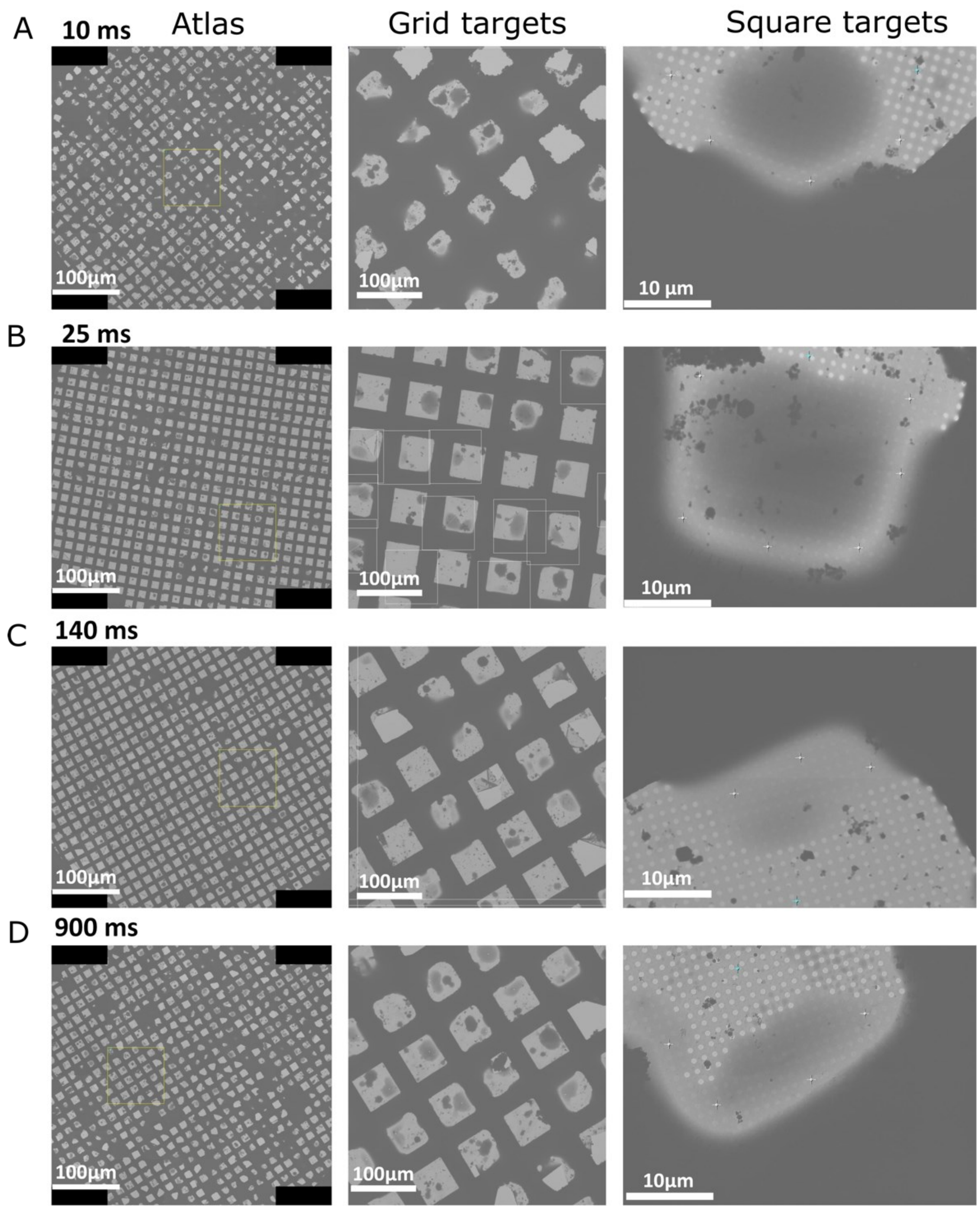


**Figure S7 | Data collection.** An atlas was screened for each grid at the different time points (left column); droplets found for square targeting (middle column); and thin ice areas on the droplet chosen for targeting of holes (right column).

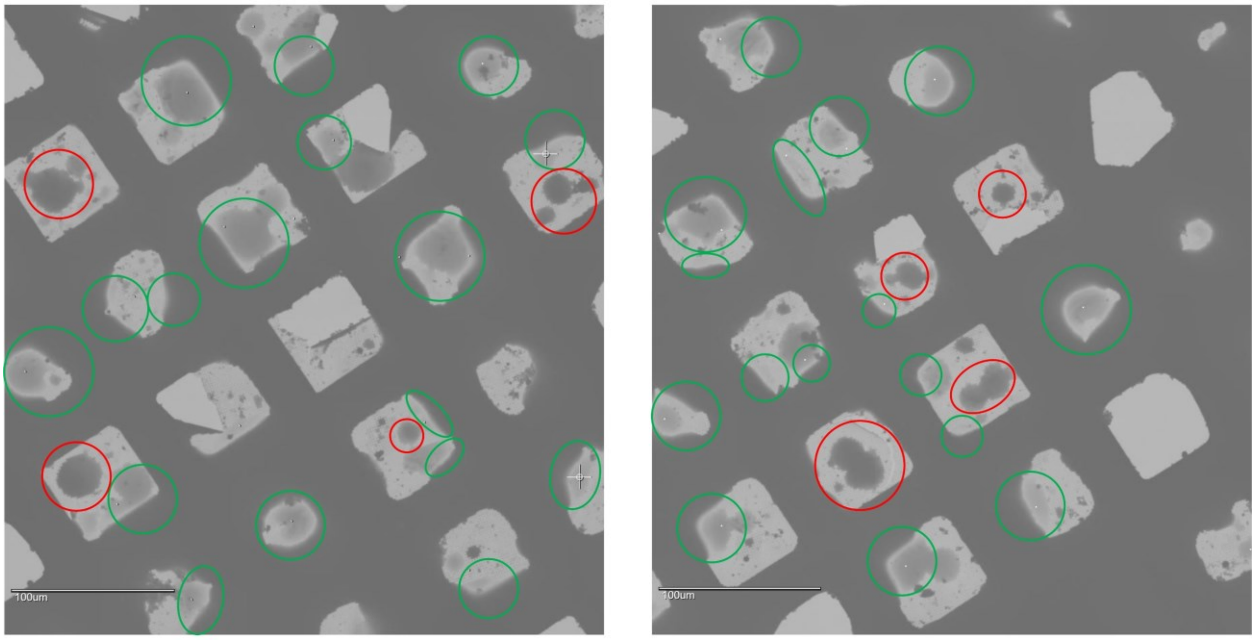


**Figure S8 | Representative examples for two types of droplets.** Those without contact with the grid bar are marked red; and those contacting the grid bar are marked green.

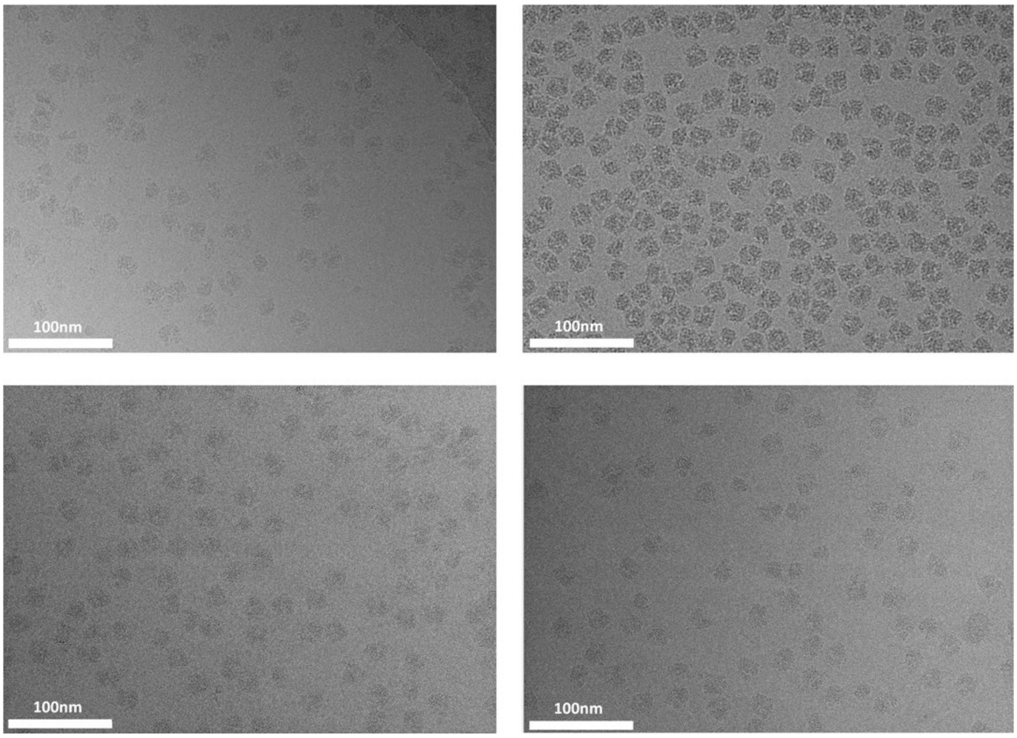


**Figure S9 | Representative micrographs from the TRCEM experiment.** High-resolution micrographs captured from the Krios cryo-electron microscope equipped with a Gatan K3 Summit camera.

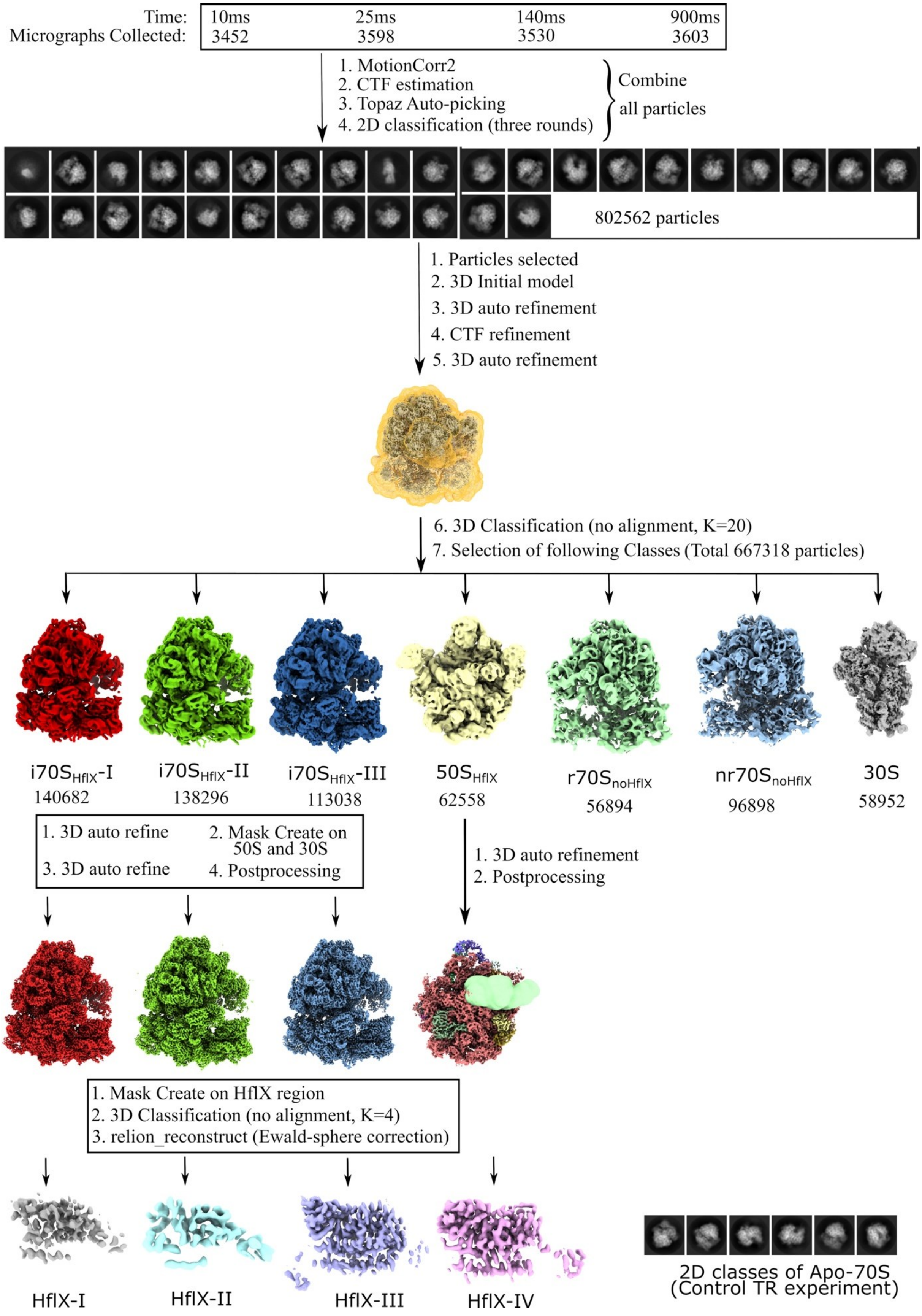

Time:
Micrographs Collected:
10ms
3452
25ms
3598
140ms
3530
900ms
3603
1. MotionCorr2
2. CTF estimation
3. Topaz Auto-picking
4. 2D classification (three rounds)
Combine all particles
802562 particles
1. Particles selected
2. 3D Initial model
3. 3D auto refinement
4. CTF refinement
5. 3D auto refinement
6. 3D Classification (no alignment, K=20)
7. Selection of following Classes (Total 667318 particles)
i70S$_{HflX}$-I
140682
i70S$_{HflX}$-II
138296
i70S$_{HflX}$-III
113038
50S$_{HflX}$
62558
r70S$_{noHflX}$
56894
nr70S$_{noHflX}$
96898
30S
58952
1. 3D auto refine
2. Mask Create on 50S and 30S
3. 3D auto refine
4. Postprocessing
1. 3D auto refinement
2. Postprocessing
1. Mask Create on HflX region
2. 3D Classification (no alignment, K=4)
3. relion_reconstruct (Ewald-sphere correction)
HflX-I
HflX-II
HflX-III
HflX-IV
2D classes of Apo-70S
(Control TR experiment)

**Figure S10 | Flow Chart of TR cryo-EM data collection and processing.** The names of the different 3D class and the corresponding numbers of particles are noted under each class.

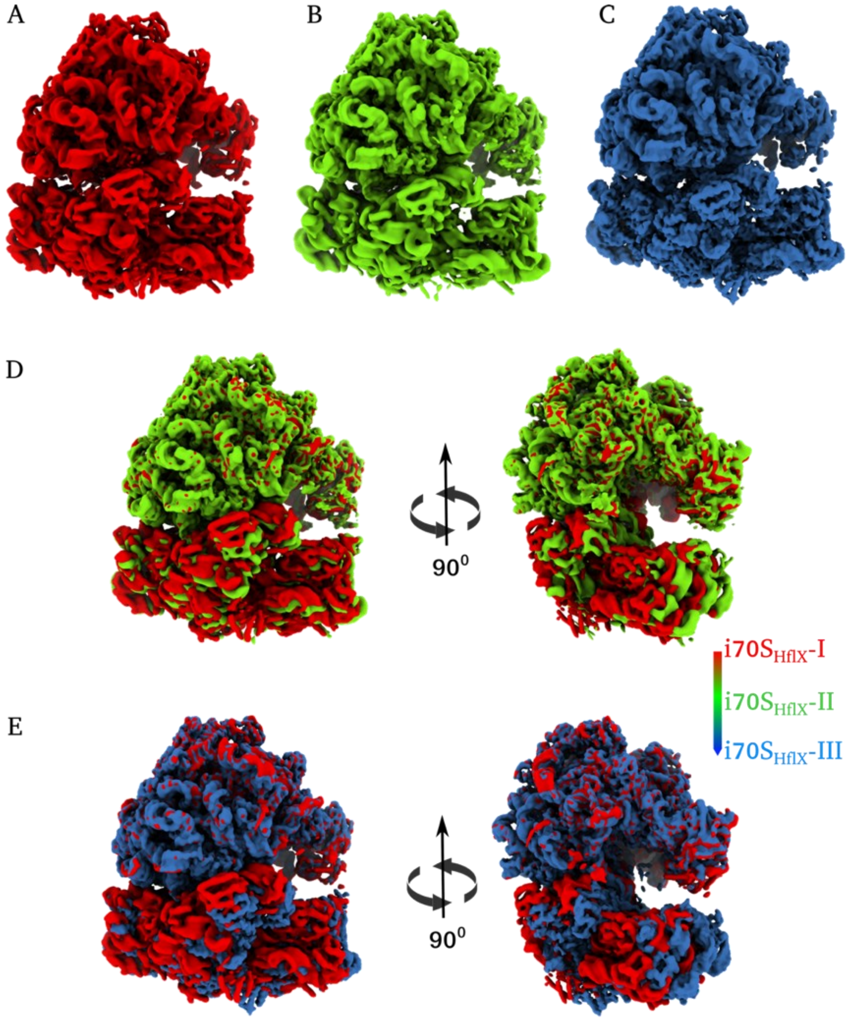


**Figure S11 | Progression of splitting of the 70S as seen in initial 3D Classes.** (**A**), (**B**), **and** (**C**), intermediate 3D classes of the $i70S_{Hflx}$-I (red), $i70S_{Hflx}$-II (green), and $i70S_{Hflx}$-III (blue), respectively. (**D**) and (**E**), superimpositions of $i70S_{Hflx}$-I (red) onto $i70S_{Hflx}$-II (green), and $i70S_{Hflx}$-I (red) onto $i70S_{Hflx}$-III (blue), respectively, to show the splitting, i.e. the downward stepwise movement of the 30S from the 50S subunit in two different views.

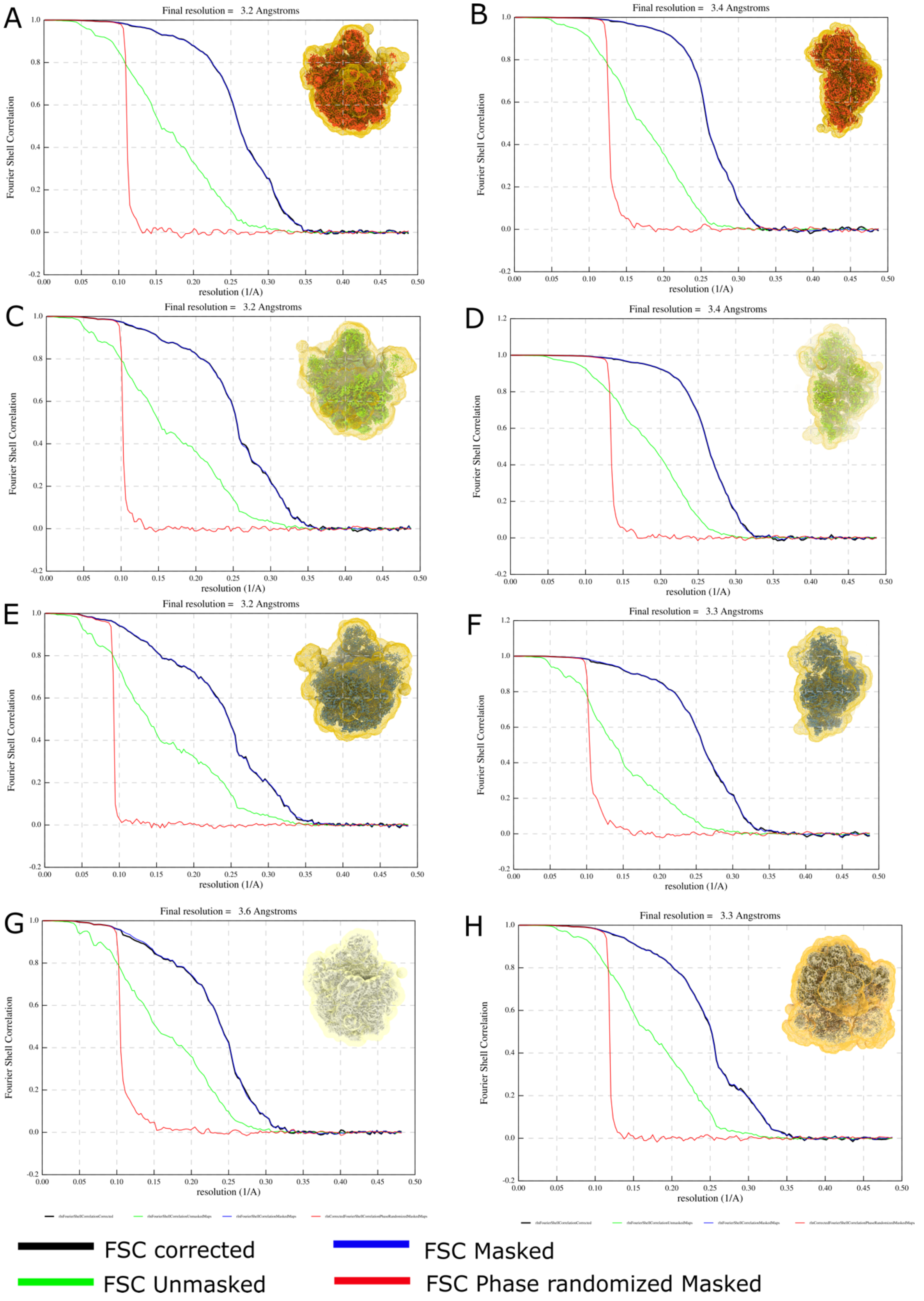


**Figure S12 | Resolution estimation via FSC.** (**A**), (**C**), (**E**), and (**G**), FSC plots and estimated resolutions for the 50S subunit of $\text{i70S}_{\text{HflX}}$-I, $\text{i70S}_{\text{HflX}}$-II, $\text{i70S}_{\text{HflX}}$-III, and for $\text{50S}_{\text{HflX}}$ itself, respectively. (**B**), (**D**), and (**F**), FSC plots and estimated resolutions for the 30S subunit of $\text{i70S}_{\text{HflX}}$-I, $\text{i70S}_{\text{HflX}}$-II, and $\text{i70S}_{\text{HflX}}$-III. The masks used and the corresponding maps are shown in the inset.

(**H**), FSC plots and resolution for the 70S consensus refinement from 802,562 particles. All representations are raw reports obtained from Relion-4.0.

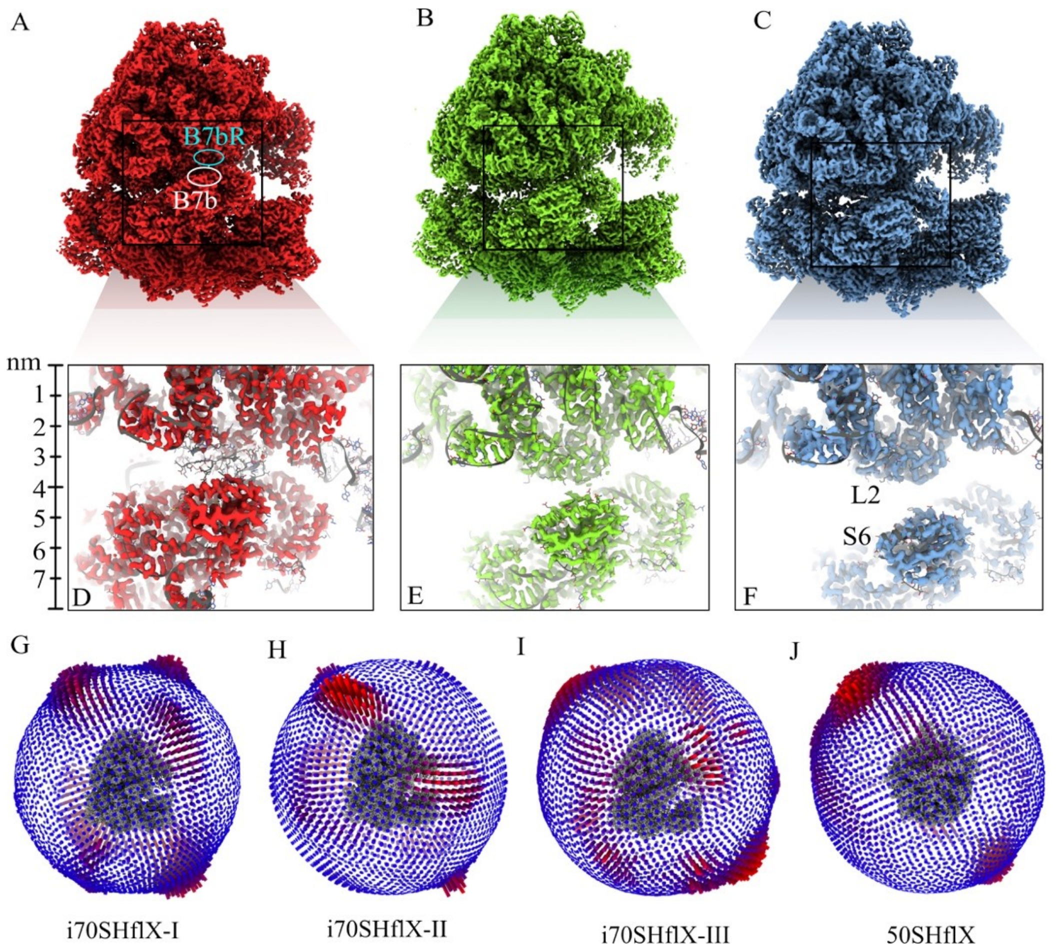


**Figure S13 |** Molecular details of subunit interface during the progressive opening of the 70S: (**A**), (**B**), and (**C**), coulomb density maps of intermediates i70SHflX-I, i70SHflX-II, and i70SHflX-III, respectively, in a view showing the separation of the subunits. All maps are aligned on the 50S subunit. (**D**), (**E**), and (**F**), zoomed views showing the time course of splitting of intersubunit bridges B7b and B7bR, and restoration of L2 from disordered (in (**D**) to its original conformation (in (**E**) and (**F**)). (**G-J**) are the angular plots of particles used.

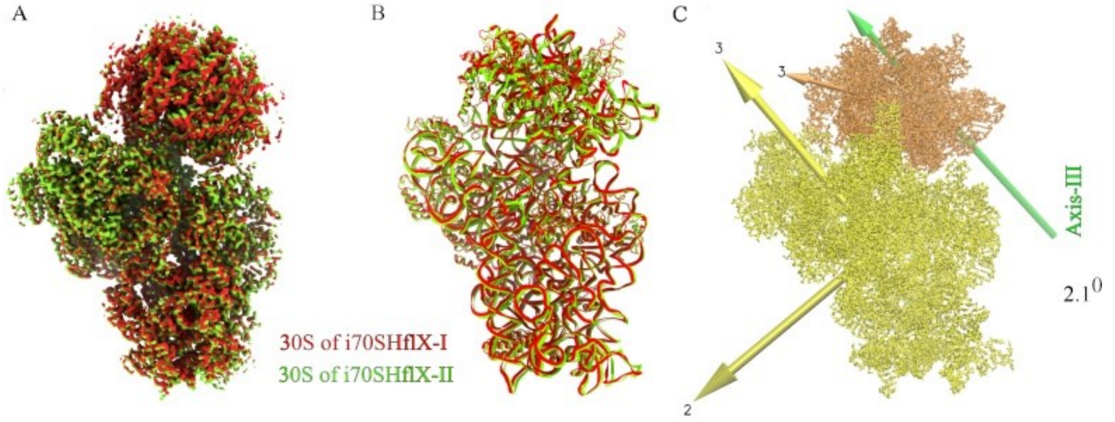


**Figure S14 | 30S head rotation during HflX-catalyzed ribosome splitting.** (**A**) and (**B**) Superimposition of maps and atomic models of the 30S from i70S$_{\mathrm{HflX}}$-I (red), and i70S$_{\mathrm{HflX}}$-II (green), respectively. (**C**) The rotation of the 30S head is calculated with respect to the 30S body. The rotation of the 30S head of i70S$_{\mathrm{HflX}}$-I by $2.1^{0}$ around hinge Axis-III (green) to adopt the conformation of the 30S subunit in i70S$_{\mathrm{HflX}}$-II.

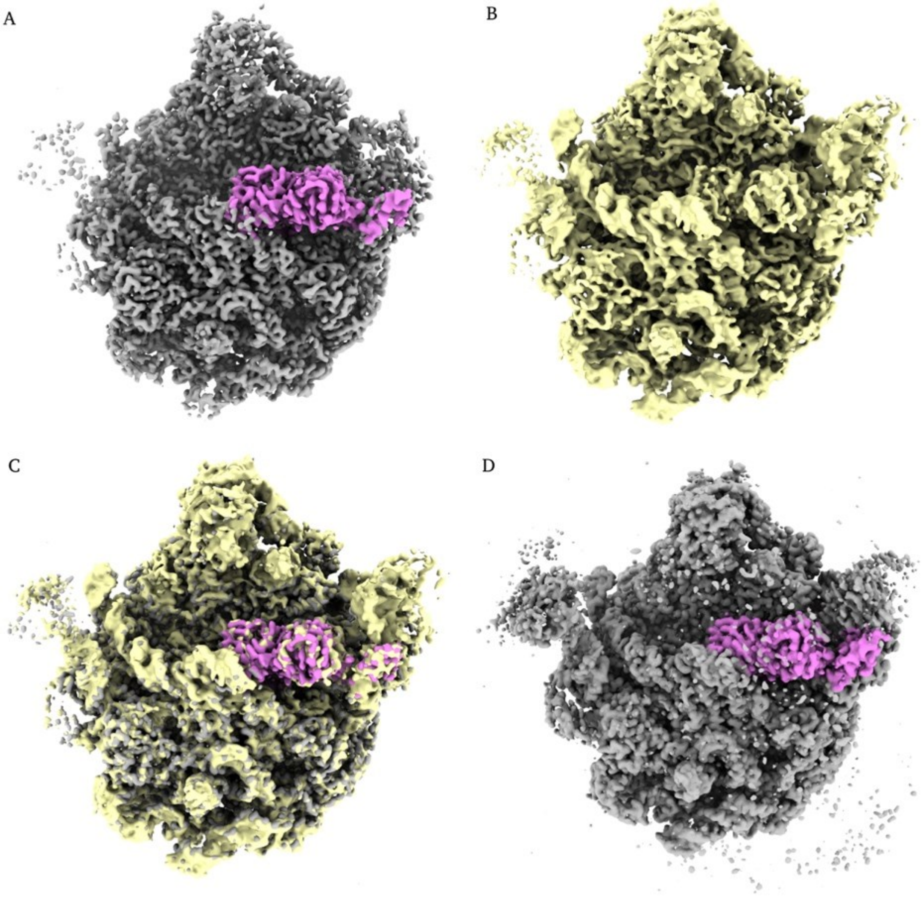


**Figure S15 | Last step of 70S splitting – comparison with literature.** (**A**) and (**B**) reconstructions of 50S$_{\mathrm{HflX}}$ (gray) and published structure of 50S-HflX-GNP-PNP (yellow, EMDB:3133). HflX in

$50S_{HflX}$ is shown in magenta and 50S in gray. (**C**) Superimposition of $50S_{HflX}$ and 50S-HflX-GNP-PNP. (**D**) view (in ChimeraX) of refined $50S_{HflX}$ structure shown in (**A**) at lower threshold level, indicating that there is no clear visible density of the 30S subunit associated with $50S_{HflX}$.

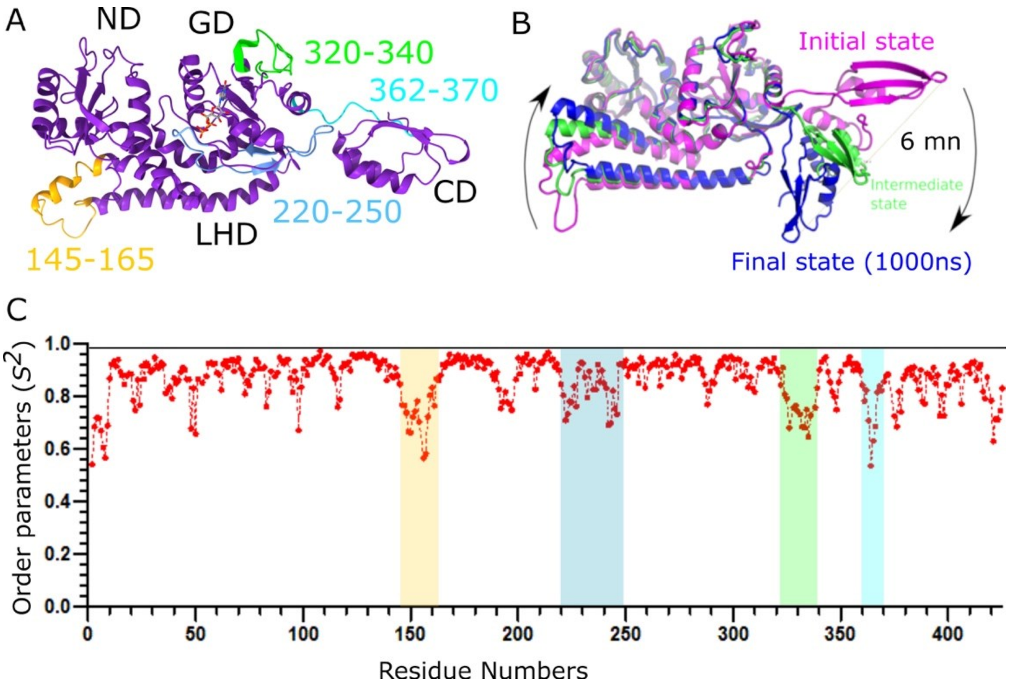


**Figure S16 | Molecular dynamics of HflX.** (**A**), Model of HflX showing its flexible region with residue numbers in various colors. (**B**), Superimposition of the three conformations extracted from 1000 ns MD simulation trajectory of free HflX. Conformations at 0 ns, 500 ns, and 1000 ns are shown in magenta, green, and blue, respectively. (**C**) Order parameters, calculated to characterize the flexible regions of HflX, and are indicated by residue zone keyed by color to HflX model in (**A**).

| Microfluidic Chips | Capillary tubing (as microreactor) inner diameter (I.D.) and length (μm/mm) | Tubing (between micromixer and microreactor) I.D. and length (μm/mm) | Tubing (between microreactor and sprayer) I.D. and length (μm/mm) | Sprayer -grid distance (mm) | Sprayer-cryogen distance (mm) | Plunging velocity (m/s) |
|---|---|---|---|---|---|---|
| Chip 1 (10 ms) | 75/5.0 | N/A | N/A | 3.5 | 10 | 1.9 |
| Chip 2 (25 ms) | 200/2.0 | 75/2.8 | 75/5.0 | 3.5 | 15 | 1.9 |
| Chip 3 (140 ms) | 200/24.0 | 75/3.0 | 75/5.4 | 3.5 | 15 | 1.9 |
| Chip 4 (900 ms) | 300/75.0 | 75/2.6 | 75/5.5 | 3.5 | 15 | 1.9 |

**Table S1 | Materials and parameters about the fabrication of the four chips used in this TR study.**

| | Control | without coating | DDM coating | $SiO_2$ coating |
|---|---|---|---|---|
| | Absorbance at 260 nm ($A_{260}$) | | | |
| Test 1 | 0.969 | 0.516 | 0.600 | 0.875 |
| Test 2 | 0.965 | 0.522 | 0.567 | 0.924 |
| Test 3 | 0.964 | 0.527 | 0.579 | 0.913 |
| Test 4 | | 0.510 | 0.605 | 0.928 |
| Test 5 | | 0.512 | 0.569 | 0.937 |
| Test 6 | | 0.519 | 0.584 | 0.925 |
| Test 7 | | | | 0.903 |
| Test 8 | | | | 0.847 |
| Averaged $A_{260}$ | 0.966 | 0.518 | 0.584 | 0.907 |
| Standard Deviation | 0.002 | 0.006 | 0.014 | 0.029 |

**Table S2 | Absorbance value at 260 nm for the sample test using chips with different coating techniques.**

| **Map and Model component** | **i70S$_{HflX}$-I** | **i70S$_{HflX}$-II** | **i70S$_{HflX}$-III** | **50S$_{HflX}$** |
|---|---|---|---|---|
| Map resolution (Å) | 50S: 3.38<br>30S: 3.19 | 50S: 3.35<br>30S: 3.2 | 50S: 3.25<br>30S: 3.2 | 50S: 3.38 |
| FSC threshold 0.143 (Å) | | | | |
| Map sharpening *B* factor (Å$^2$) | 30S: -43.89<br>50S: -56.43 | 30S: -70.01<br>50S: -56.99 | 30S: -49.97<br>50S: -63.22 | 50S: -73.64 |
| Model composition (residues) | | | | |
| Protein | 5922 | 5922 | 5922 | 3799 |
| Nucleotide | 4559 | 4559 | 4559 | 3020 |
| Mg$^{2+}$ ions | | | 1 | 1 |
| GTP/GDP-Pi | | | 1 | 1 |
| waters | | | | 2 |
| *B* factors (Å$^2$) | | | | |
| RNA | 55.05 | 80.74 | 54.47 | 95.02 |
| protein | 43.84 | 77.83 | 45.84 | 78.62 |
| R.m.s. deviations from ideal values | | | | |
| Bond (Å) | 0.008 | 0.006 | 0.008 | 0.006 |
| Angle (°) | 1.009 | 0. 950 | 1.052 | 0.965 |
| Molprobity score | 1.56 | 1.69 | 1.59 | 1.85 |
| Clash score | 4.56 | 5.22 | 5.44 | 7.51 |
| Ramachandran plot (%) | | | | |
| Favored (%) | 95.25 | 93.79 | 96.48 | 93.20 |
| Allowed (%) | 4.68 | 6.16 | 3.47 | 6.72 |
| Outliers (%) | 0.07 | 0.05 | 0.05 | 0.08 |
| Rotamer outliers (%) | 0.68 | 0.74 | 0.80 | 0.84 |
| CB outliers (%) | 0.06 | 0.02 | 0.07 | 0.06 |
| RNA validation | | | | |
| Good_Sugar Puckers(%) | 99.99 | 99.99 | 99.63 | 99.99 |
| Good_backbone_conformation(%) | 99.74 | 99.99 | 99.98 | 99.98 |
| Average suiteness | 0.503 | 0.518 | 0.517 | 0.507 |

**Table S3 | Model refinement statistics.**
